# Role of Ring6 in the function of the *E. coli* MCE protein LetB

**DOI:** 10.1101/2021.09.30.462657

**Authors:** Casey Vieni, Nicolas Coudray, Georgia L. Isom, Gira Bhabha, Damian C. Ekiert

**Affiliations:** Skirball Institute of Biomolecular Medicine and Department of Cell Biology, New York University School of Medicine, New York, NY; Department of Microbiology, New York University School of Medicine, New York, NY; Applied Bioinformatics Laboratories, New York University School of Medicine, New York, NY, USA

**Keywords:** cryo-EM, lipid transport, bacterial cell envelope, microbiology, LetB, MCE

## Abstract

LetB is a tunnel-forming protein found in the cell envelope of some double-membraned bacteria, and is thought to be important for the transport of lipids between the inner and outer membranes. In *Escherichia coli* the LetB tunnel is formed from a stack of seven rings (Ring1 - Ring7), in which each ring is composed of a homo-hexameric assembly of MCE domains. The primary sequence of each MCE domain of the LetB protein is substantially divergent from the others, making each MCE ring unique in nature. The role of each MCE domain and how it contributes to the function of LetB is not well understood. Here we probed the importance of each MCE ring for the function of LetB, using a combination of bacterial growth assays and cryo-EM. Surprisingly, we find that ΔRing3 and ΔRing6 mutants, in which Ring3 and Ring6 have been deleted, confer increased resistance to membrane perturbing agents. Specific mutations in the pore-lining loops of Ring6 similarly confer increased resistance. A cryo-EM structure of the ΔRing6 mutant shows that despite the absence of Ring6, which leads to a shorter assembly, the overall architecture is maintained, highlighting the modular nature of MCE proteins. Previous work has shown that Ring6 is dynamic and in its closed state, may restrict the passage of substrate through the tunnel. Our work suggests that removal of Ring6 may relieve this restriction. The deletion of Ring6 combined with mutations in the pore-lining loops leads to a model for the tunnel gating mechanism of LetB. Together, these results provide insight into the functional roles of individual MCE domains and pore-lining loops in the LetB protein.

**Highlights:**

- Deleting MCE domains 3 or 6 from LetB confers increased resistance to membrane-perturbing agents
- Cryo-EM structure of ΔRing6 LetB mutant highlights the modular nature of MCE domains
- Mutations along LetB pore-lining loops modulate detergent resistance

## Introduction

The Gram-negative bacterial cell envelope is composed of an inner membrane (IM) and an outer membrane (OM), separated by an aqueous periplasm containing the peptidoglycan cell wall. The OM protects the cell from extracellular stress, but also poses unique challenges for the transport of molecules across the cell envelope. Macromolecular machines, such as efflux pumps and lipid transporters, have evolved to transport specific hydrophobic substrates across the hydrophilic periplasmic space that separates the IM from the OM (Konovalova and Silhavy, 2015; Shi et al., 2019; Wang et al., 2017; Wong et al., 2019). For example, the lipopolysaccharide (LPS) transport system utilizes an IM ABC transporter (LptBFG) to translocate LPS along a periplasmic bridge (LptCA) to an OM β-barrel protein (LptDE), which then inserts LPS into the outer leaflet of the OM (Chng et al., 2010; Dong et al., 2014; Li et al., 2019; Owens et al., 2019; Qiao et al., 2014; Sperandeo et al., 2007). Other systems, such as the multidrug efflux pumps MacAB-TolC and AcrAB-TolC, form an enclosed tunnel across the periplasm for substrate transport. In recent years, the MCE family of transporters (which contain **M**ammalian **C**ell **E**ntry protein domains) (Arruda et al., 1993; Cantrell et al., 2013; Klepp et al., 2012; Marjanovic et al., 2011) has been implicated in the transport of lipids (Ekiert et al., 2017; Isom et al., 2017; Malinverni and Silhavy, 2009; Nakayama and Zhang-Akiyama, 2017) and other hydrophobic molecules (Cantrell et al., 2013; Kim et al., 1998; Marjanovic et al., 2011; Mohn et al., 2008; Pandey and Sassetti, 2008) across the periplasm in diderm bacteria. In particular, two MCE proteins from *E. coli*, LetB and PqiB, have been shown to form tunnels capable of spanning the periplasm, though their precise transport mechanism remains poorly understood.

LetB (previously known as YebT, and homologous to MAM-7 from *Vibrio parahemolyticus*; (Krachler et al., 2011; Krachler and Orth, 2011) is encoded in an operon together with the integral IM protein *letA* (formerly *yebS*). LetB belongs to the MCE protein family and forms a **l**ipophilic **e**nvelope spanning **t**unnel that is long enough to span the cell envelope, from the IM to the OM (Ekiert et al., 2017; Isom et al., 2020; Liu et al., 2020). It is therefore poised to facilitate transport of lipids or other hydrophobic molecules across the cell envelope. The LetB polypeptide consists of seven tandem MCE domains, and cryo-EM structures have revealed that these seven domains are arranged in a linear fashion, like beads on a string in a single LetB protomer. Six LetB monomers associate with each other to form a homohexameric tunnel in which a given MCE domain interacts with the equivalent MCE domain from a neighboring subunit. Thus, MCE1 from each protomer comes together to form a hexameric ring, followed by MCE2, MCE3, MCE4, MCE5, MCE6 and MCE7 (Fig. 1A) (Isom et al., 2020; Liu et al., 2020). The length of the LetB tunnel is determined by the number of MCE domains encoded in the primary sequence, with each MCE domain resulting in one ring. LetB variants have been engineered with a range of tunnel lengths by deleting one or more MCE domains, and naturally occurring MCE proteins can encode from one to eight MCE domains in their primary sequence, resulting in proteins made of one to eight stacked rings (Isom et al., 2020). This suggests that MCE proteins are modular, consisting of MCE domain modules. The functional role of each MCE module is not well understood.

**Figure 1.**
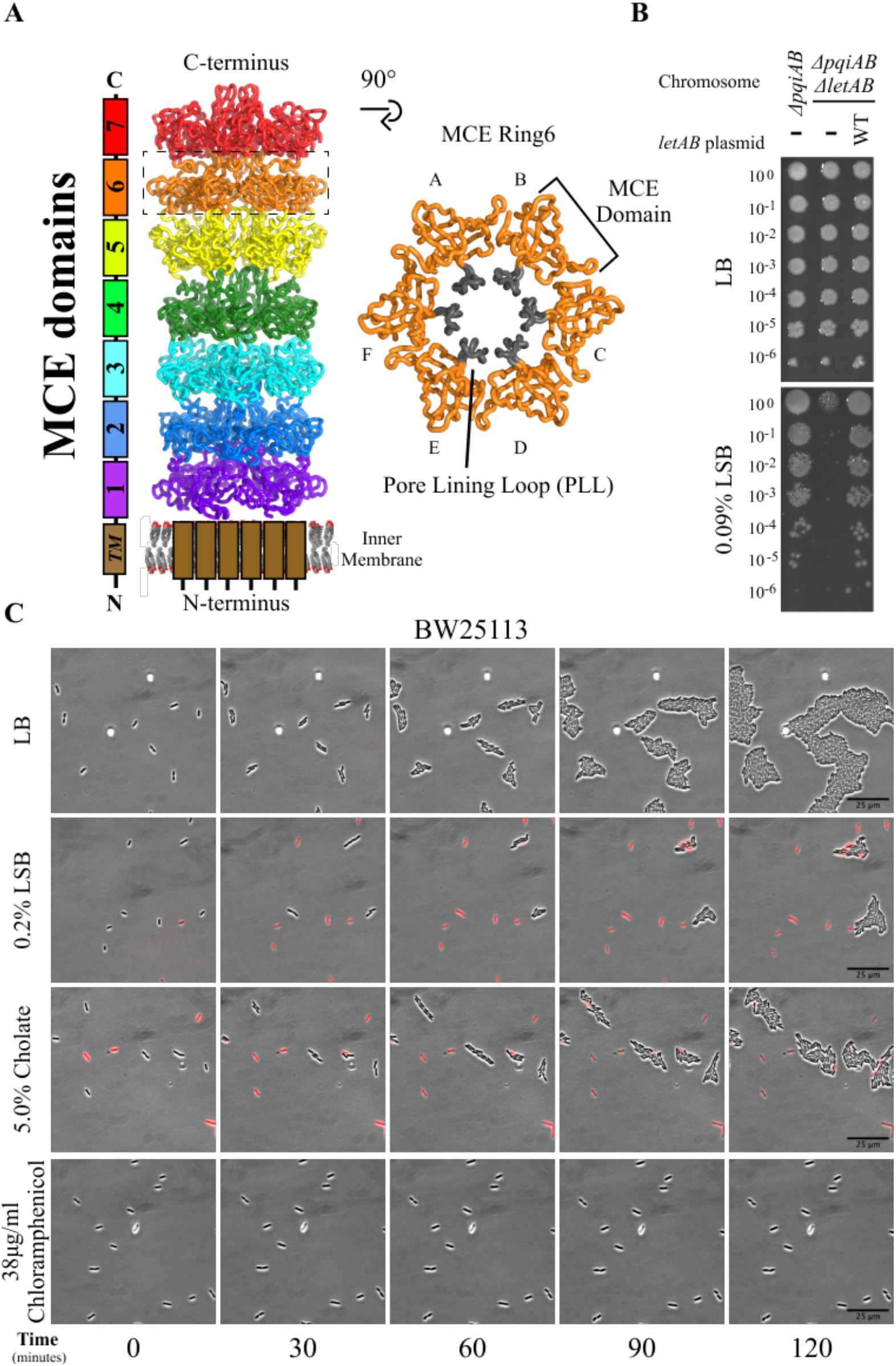
Mechanism of action of LSB and cholate in *E. coli*. **(A)** Side-view of the structure of WT LetB (PDB 6V0D), and a top view of the homohexameric ring formed by MCE6. The MCE domains 1-7 are shown as different colors. **(B)** The *ΔpqiAB ΔletAB* strain grows poorly in the presence of LSB compared to the parent *ΔpqiAB* strain, and can be complemented by the expression of WT *letAB* from a plasmid. 10-fold serial dilutions of the specified cultures were spotted on LB agar containing any additives as indicated, and incubated overnight. **(C)** Representative time-lapse image series showing growth of *E. coli* BW25113 on LB, 0.2% LSB, 5.0% cholate, or 38 μg/ml chloramphenicol. Displayed images are a merge of the phase contrast and the PI channels. Unmerged images are shown in Fig. S2.

The interior surface of the LetB tunnel is formed by the pore-facing regions of each MCE domain, and is largely hydrophobic. Each MCE domain contains a pore-lining loop (PLL) that emerges from the domain and forms the lining of the pore (Fig. 1A). The PLLs are dynamic, and the diameter of the LetB pore is modulated by conformational changes in the MCE rings and PLLs (Isom et al., 2020). The modulation of the LetB pore diameter may be important for passage of substrate through the pore, and therefore MCE rings may play a role in regulating the transport of substrates through the LetB pore. Here, we performed a structure-function analysis of LetB, with the goal of addressing the functional roles of different MCE domains in the protein.

## Results

### LSB and Cholate are Bactericidal

Deletion of *letAB* renders *E. coli* modestly more sensitive to cholate and deoxycholate (Isom et al., 2020), as well as bile salts (Fig. S1, A, B, E; Note: ΔRing6 mutant will be discussed later in the manuscript). *LetAB* mutants are also mildly more resistant to the antibiotics vancomycin (Isom et al., 2017) (Fig. S1C) and rifampicin (Fig S1D). *LetAB* mutants are also sensitive to lauryl sulfobetaine (LSB) when combined with mutations in a related MCE transport system called *pqi*, resulting in a 2-3 log reduction in growth of a *pqiAB letAB* double mutant compared with the parental *pqiAB* strain (Isom et al., 2020, 2017) (Fig. 1B). Primary bile salts, such as cholate, and secondary bile salts, such as deoxycholate, emulsifying agents that are synthesized in the liver and intestinal tracts of vertebrates, and also serve as toxic detergents to enteric bacteria by interfering with biological membranes (Bortolini et al., 1997; Gunn, 2000; Helenius and Simons, 1975; Nikaido, 2003). LSB is a synthetic, zwitterionic surfactant with antimicrobial properties used in industrial applications (Chen et al., 2011; He et al., 2009; Wieczorek et al., 2016; Zhang et al., 2018). The sensitivity to LSB and bile salts observed upon deletion of *letAB* is consistent with a mild defect in bacterial outer membrane homeostasis (Burman et al., 1972; Ma et al., 1995; Nikaido, 2003; Poole, 2004; Ruiz et al., 2006; Thanassi et al., 1997; Wang, 2002).

While these data show that cells lacking functional LetAB are sensitive to various membrane-perturbing agents, the mechanism of action of these compounds is not well understood. We chose to focus on LSB, since it has been previously used to investigate LetAB function, and cholate, as a representative bile salt. To assess whether LSB and cholate are bactericidal or bacteriostatic, we used live-cell fluorescence microscopy (see methods) to assess how LSB and cholate affect *E. coli* growth and survival on solid media. We stained cells with the membrane-impermeable dye propidium iodide (PI), which cannot cross the cell envelope when it is intact but can penetrate compromised membranes of dead cells (Boulos et al., 1999). Thus, cells that are stained with PI (PI+) are likely non-viable, while cells that are not stained with PI (PI-) are viable. In the absence of LSB, PI+ cells were very rare (Fig. 1C, Fig. S2). After 60 minutes in the presence of 0.2% LSB, 25 ± 30% cells become PI+, and only a few cells survive and go on to form microcolonies (Fig. 1C, Fig. S2). Similarly, *E. coli* grown in the presence of 5.0% cholate resulted in 20 ± 16% cells rapidly becoming PI+ within the first 3-5 minutes, and only a few cells survive to form colonies (Fig. 1C, Fig. S2). In contrast, *E. coli* grown in the presence of chloramphenicol, a known bacteriostatic agent (Brock, 1961; Fassin et al., 1955; Wisseman et al., 1954), remain predominantly PI-over the two hour time course and none of the cells go on to form colonies (Fig 1C, Fig. S2). This suggests that LSB and cholate are bactericidal rather than bacteriostatic, and may act by disrupting cellular membranes.

### ΔRing6 and ΔRing3 LetB mutants are more resistant to LSB and cholate

LetB assembles into an elongated barrel formed by a stack of seven rings, in which each ring is a homohexamer of a given MCE domain (Fig. 1A) (Ekiert et al., 2017; Isom et al., 2020; Liu et al., 2020). The LetB protein can be engineered to have fewer than seven rings by deleting individual MCE domains from the gene (Fig. 2A). Previous work demonstrated that the WT, ΔRing6 mutant (deletion of Ring6) or a ΔRing5-6 mutant, which result in LetB proteins that are seven, six, or five rings long respectively, conferred LSB resistance in a *pqiAB letAB* knockout background. However, mutants encoding LetB proteins with a total length of four rings or less were not resistant to LSB (Isom et al., 2020). These data suggested that the overall length of LetB contributes to its function, but the role of specific rings, and whether the rings are functionally equivalent to each other, remains an open question. Due to the modular organization of LetB, we reasoned that we could assess the relative functional contribution of MCE rings by deleting each ring individually and assessing the ability of the resulting proteins to function in cells (Fig. 2A). We designed single ring deletion mutants (see methods for details of construct design), and assessed their ability to restore growth on LSB when introduced into a *ΔpqiAB ΔletAB* strain. We used the LSB phenotype for our genetic complementation assays as it is the most robust phenotype for *letB* (Fig. 1B). Complementing with a plasmid containing wild type *letAB* can restore growth to the level of the parent, *ΔpqiAB* strain (Fig. 1B). ΔRing1, ΔRing2, ΔRing5, or ΔRing7 mutants failed to restore growth on LSB, while the ΔRing4 mutant restored growth on LSB to a similar level as WT LetAB. Strikingly, ΔRing3 or ΔRing6 mutants conferred an even higher level of resistance to LSB than the WT (Fig. 2B, Fig. S3A). A *ΔletAB* strain complemented with a ΔRing6 mutant also shows substantially increased resistance to cholate, compared with the WT (Fig. 2C). The ΔRing3 and ΔRing4 mutants also show increased resistance to cholate compared with WT, but to a lesser degree than the ΔRing6 mutant. We found that our phenotypic assay on LSB more clearly shows differences between the parental and knockout strains, and hence can be better used to differentiate strains that complement vs. strains that do not complement. The cholate condition more clearly highlights increased resistance. Based upon their locations at either end of the LetB tunnel, Ring1 and Ring7 have been proposed to interact with the inner and outer membrane and likely other membrane proteins; therefore it is expected that deleting these rings may disrupt the ability of LetB to transfer lipids or other substrates to or from the IM and OM. In contrast, the remaining rings (Ring2 - Ring6) stack along the middle of the tunnel and contribute to its overall length. The observation that removal of some rings results in functional LetB while removal of others does not, may indicate that some rings have specialized functions.

**Figure 2.**
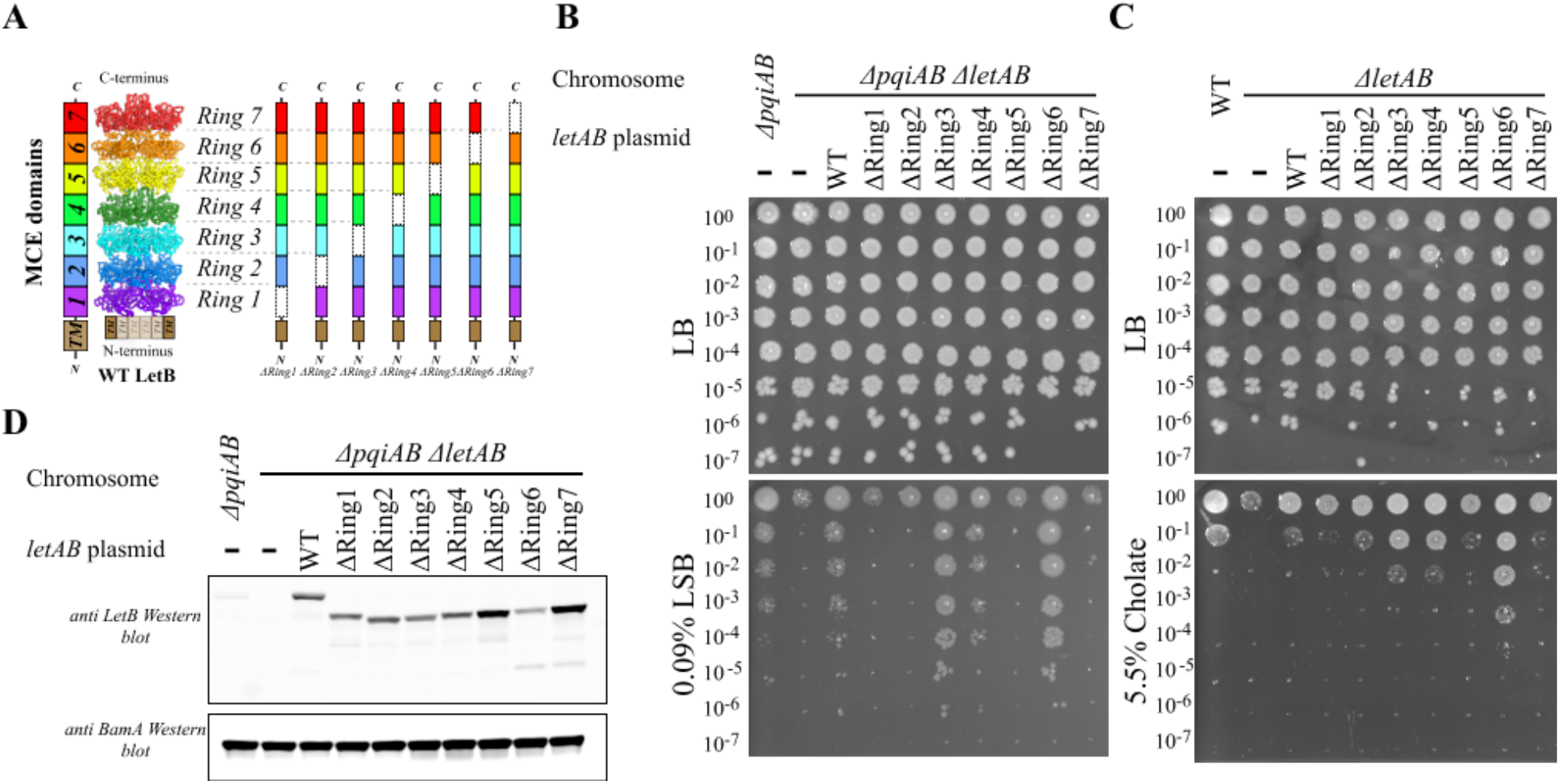
ΔRing6 and ΔRing3 mutants exhibit increased resistance to LSB and cholate. **(A)** Schematic diagram of LetB domain organization and ring deletion mutants. Deleted MCE domains are shown as dashed rectangles. **(B and C)** Cellular assay for the function of LetB ring deletion mutants based on LSB resistance **(B)** or cholate resistance **(C)**. 10-fold serial dilutions of the specified cultures were spotted on LB agar containing any additives as indicated, and incubated overnight. **(D)** Anti-LetB western blot to compare cellular levels of WT LetB and ring deletion mutants. Membrane-enriched fractions were prepared from all strains used in the complementation experiments and tested for expression of LetB using a rabbit polyclonal antibody against LetB. As a control, a rabbit polyclonal antibody against BamA was used. BamA serves as an OM marker to assess cell envelope homeostasis.

Based on our observation that ΔRing3 and ΔRing6 mutants show increased resistance to LSB, we wondered whether this phenotype was due to increased expression levels of these mutants. To determine whether this is the case, we assessed the levels of LetB protein in the complemented strains by Western blotting using an antibody against LetB (Fig. 2D; Fig. S3B). All complemented strains express LetB at a higher level than endogenous LetB in the parent strain. ΔRing3 and ΔRing6 do not show an increased level of expression relative to other mutants or the WT complemented strain. Therefore, the increased LSB resistance of the ΔRing3 and ΔRing6 mutants cannot readily be explained by increased LetB expression levels and may instead be due to specialized functions of these rings.

### Cryo-EM Structure of ΔRing6 LetB

The ΔRing6 mutant exhibited the most resistance to LSB in our phenotypic growth assays. To assess if any structural changes may explain why deleting Ring6 leads to increased LSB resistance, we used single particle cryo-EM to determine the structure of the periplasmic domain of the ΔRing6 mutant (residues 43–632, 747-877) (Fig. 3A). We used a similar data processing pipeline as previously reported for WT LetB (Isom et al., 2020) in which particle picking and 2D classification were followed by iterative 3D classification, signal subtraction, and focused refinement (Fig. S4). We reconstructed two density maps with resolutions of 3.2 Å and 3.6 Å, allowing us to build complete structural models of ΔRing6 LetB (see Fig. S5A, S5B, S5C, S5D and methods for further details regarding model building). The two resulting models differ primarily in the open and closed configuration of MCE Ring1. Coordinates and maps are available for immediate download as supplementary files associated with this preprint, and will be publicly released from the PDB/EMDB as soon as the entries are processed.

**Figure 3.**
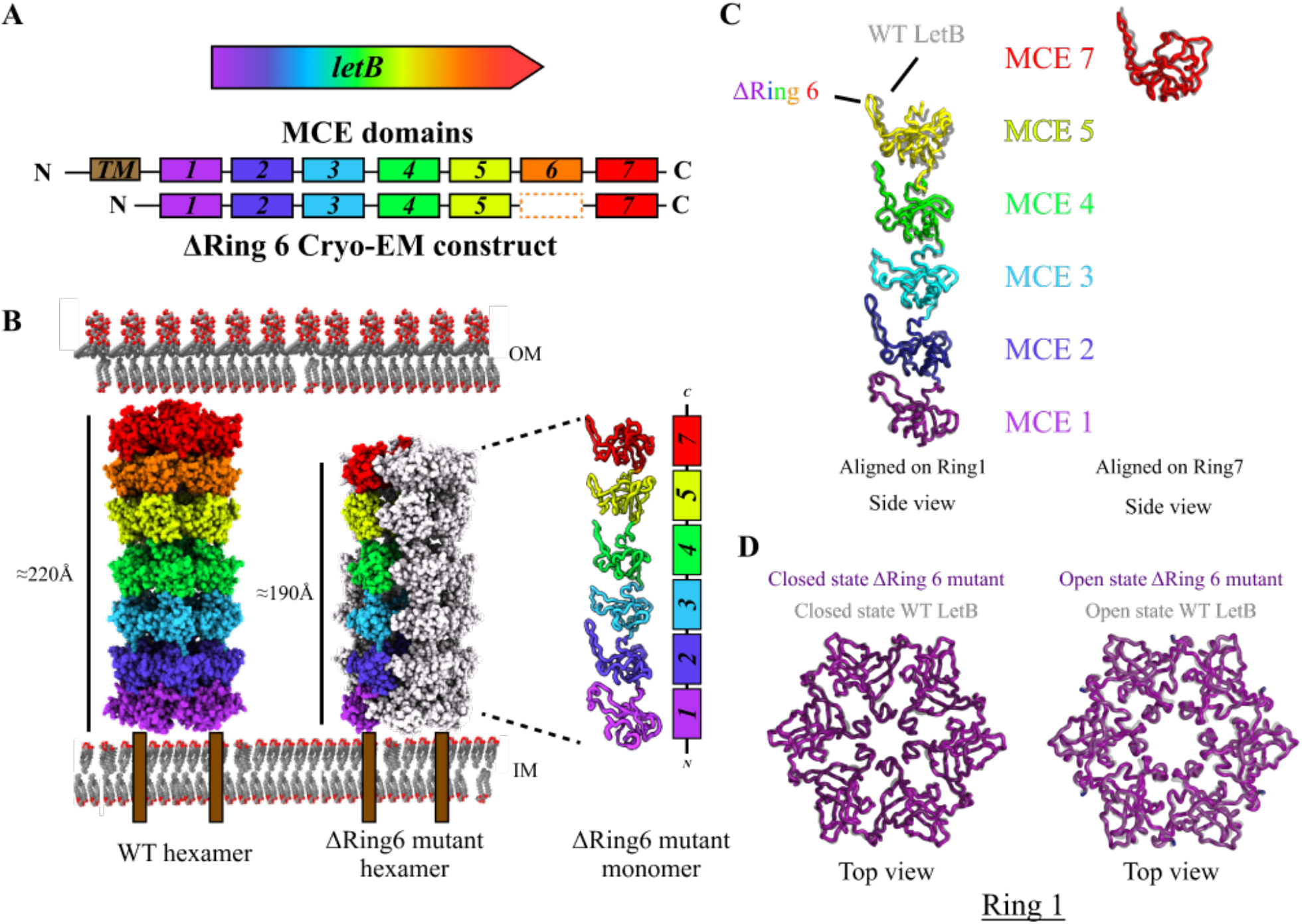
Cryo-EM structure of ΔRing6 LetB mutant. **(A)** Schematic diagram of WT LetB vs. the ΔRing6 construct used for structure determination by single particle cryo-EM. **(B)** Comparison of WT (PDB ID: 6V0D) and ΔRing6 LetB structures. In the ΔRing6 LetB structure, a single protomer is colored. The ΔRing6 mutant is about 30Å shorter, corresponding to the removal of MCE6. **(C)** Overlay of WT (PDB ID: 6V0D) and ΔRing6 structures, aligned on the ring as indicated. One protomer is shown. **(D)** Comparison of closed and open states of Ring1, as observed in the WT (PDB ID: 6V0C and 6V0J, respectively) and ΔRing6 structures (model #1 and model #2, respectively).

As expected, ΔRing6 LetB is shorter than WT LetB by one ring, resulting in an overall length of 190 Å, compared with 220 Å for WT LetB (Fig. 3B). Despite removing an entire ring, the overall arrangement of domains in the ΔRing6 mutant is very similar to WT LetB (Fig. 3C) (Isom et al., 2020; Liu et al., 2020), highlighting the modular nature of the LetB protein assembly. Each protomer of ΔRing6 consists of six sequential MCE domains with six protomers homo-hexamerizing to form a tunnel (Fig. 3B, C, D). Each MCE domain of ΔRing6 is nearly identical in structure to its counterpart in the WT LetB structure (average rmsd 0.61 Å, range: 0.34 Å for MCE2 between 6V0D and ΔRing6 to 1.2 Å for MCE5 between 6V0D and ΔRing 6). In the ΔRing6 protein, a new interface is generated between Ring5 and Ring7, but the positioning of MCE domains in Ring5 and Ring7 is very similar to WT LetB.

In previous analyses of the WT structure, two main conformational states were observed, a “closed” state and an “open” state. For WT LetB, Ring1, and separately, Rings5-7, were observed in closed and open states. In the Rings5-7 module, Ring6 clearly adopted one state in which the pore was open (PDB 6V0D), and a second state in which the pore was closed (PDB 6V0C). When closed, Ring6 forms a constriction point in the tunnel, and would block substrate from passing through, leading to a model in which Ring6 may gate transport through the tunnel. We asked whether the conformational landscape of ΔRing6 LetB was similar or different. However, despite extensive signal subtraction and 3D classification, we were unable to observe any alternate conformations for the Ring5+7 module, suggesting that either removing Ring6 has altered the dynamic landscape or we are unable to separate the different conformational states. In WT LetB, Ring6 undergoes the largest conformational change, possibly driving the 3D classification and allowing us to tease apart different conformational states. The absence of Ring6 may be one reason that we are unable to classify conformational states for the Rings5-7 module in the ΔRing6 mutant, if they exist. For Ring1 in the ΔRing6 mutant, we observed two conformational states, very similar to the closed (Map/Model #1) and open (Map/Model #2) states for Ring1 in the WT protein (Fig. 3D). In addition, multi-body refinement in RELION (Zivanov et al., 2018) revealed classes with potential rotation of Rings5+7 and Ring1 relative to Rings2-4 similar to what was reported for WT LetB (Isom et al., 2020). As the cryo-EM structure shows that the ΔRing6 mutant differs from the WT LetB structure primarily in the absence of Ring6, while the overall architecture is well preserved, the increased LSB resistance may be due to the loss a specialized function, such as gating substrate transport when Ring6 is in the closed conformation.

### Mutation of conserved hydrophobic residues in LetB tunnel abolish function

The opening and closing of Ring6 in WT LetB involves the coordinated rotation of the MCE domains that form Ring6, as well as movements of the PLL which line the tunnel. The PLLs both create the tunnel lining in the open state, as well as physically occlude the tunnel in the closed state, and were previously shown to be essential for LetB function (Isom et al., 2020). Each PLL consists of two main parts: 1) a hairpin loop running roughly parallel to the tunnel length, and 2) a short helix near the base of the hairpin. Previous work found that each PLL in LetB contains a conserved hydrophobic motif ΦxxΦΦ (Φ denotes a hydrophobic residue; x denotes any residue), which maps to the short helix of each PLL and positions the three conserved hydrophobic residues towards the interior of the LetB tunnel (Fig. 4A) (Isom et al., 2020). Single amino acid substitutions in the hydrophobic motifs in PLL2 and PLL3 completely abolish LetB function, and we sought to assess whether the conserved motifs in PLLs 1, 4, 5, 6, and 7 were also important for LetB function. To this end, we constructed a panel of single mutants in which each of the 3 conserved hydrophobic residues in each of the 7 PLLs was mutated to Asn (see methods). Of the 21 mutations constructed, 20 mutants failed to complement the growth of a *ΔpqiAB ΔletAB* strain in the presence of LSB (Fig. 4B). The only exception was A598N, which was able to partially restore growth. Thus, despite a large number of residues facing towards the tunnel interior, single mutations in these conserved hydrophobic motifs of LetB are sufficient to render the protein completely non-functional. As several of these mutants have been shown to still fold and assemble normally (Isom et al., 2020), this suggests that these motifs are key functional sites in LetB.

**Figure 4.**
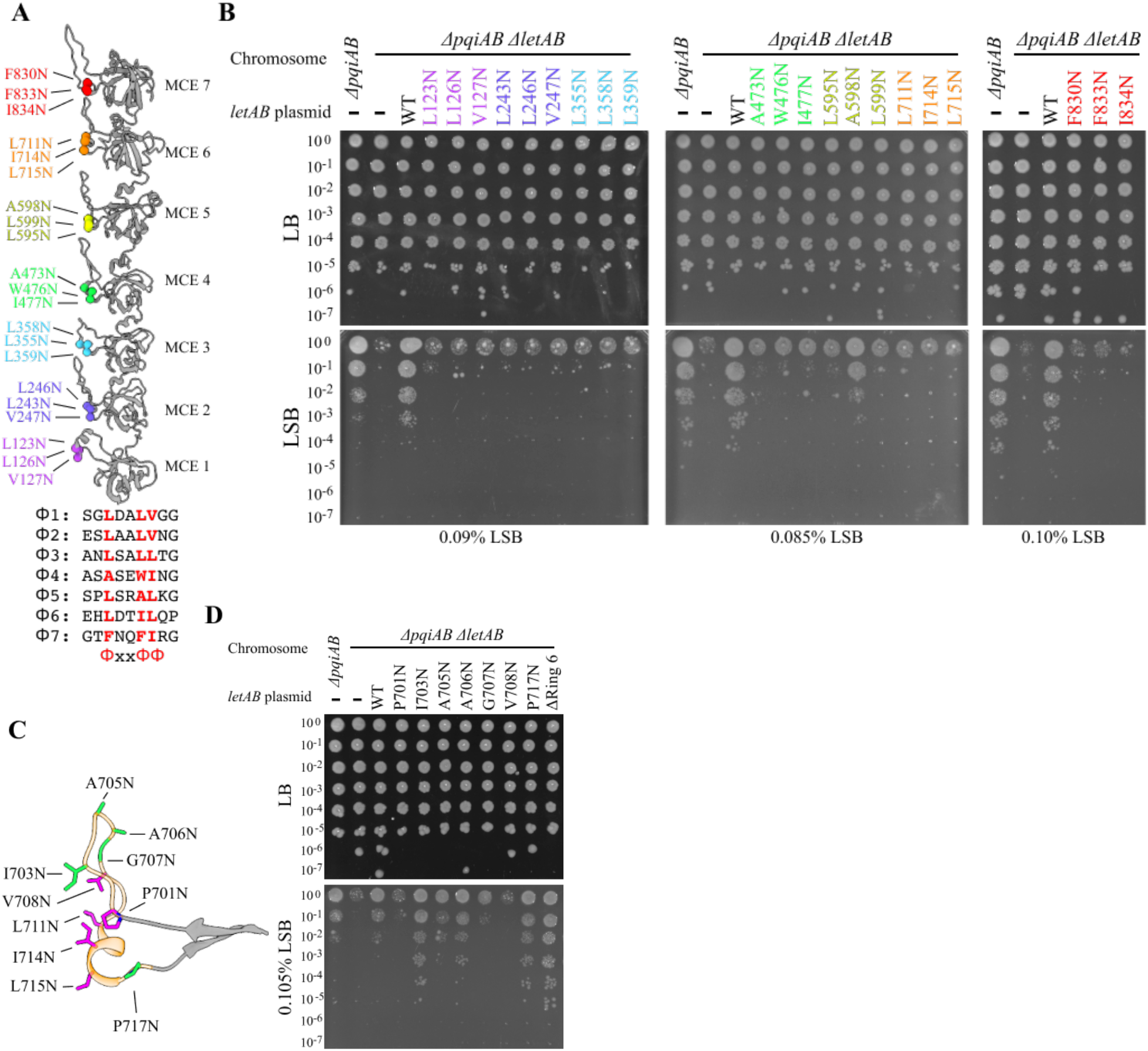
Phenotypes of point mutants in the hydrophobic motif of each MCE domain. **(A)** Residues of hydrophobic motifs are shown as colored spheres mapped on a LetB protomer. Sequence alignment of LetB hydrophobic motifs is shown below the model. **(B)** Cellular assay for the function of LetB hydrophobic motif mutants based on LSB resistance. 10-fold serial dilutions of the specified cultures were spotted on LB agar containing any additives as indicated, and incubated overnight. All mutations except for A598N attenuate LetB function and result in growth indistinguishable from the *ΔpqiAB ΔletAB* strain. **(C)** Hydrophobic residues along PLL6 are shown as sticks. Mutations resulting in strains that do not restore growth on LSB in a *ΔpqiAB ΔletAB* background are shown in magenta and mutations resulting in strains that restore growth on LSB in a *ΔpqiAB ΔletAB* background are shown in green. **(D)** Cellular assay for the function of LetB PLL6 scanning mutants based on LSB resistance. 10-fold serial dilutions of the specified cultures were spotted on LB agar containing any additives as indicated, and incubated overnight.

### Mutations in PLL6 can increase LSB resistance

In addition to the residues that make up the hydrophobic motif, several other hydrophobic side chains in each PLL are positioned towards the lumen of the tunnel, creating a generally hydrophobic tunnel lining. To further explore whether mutating any hydrophobic residue along a PLL would attenuate LetB function, we carried out scanning mutagenesis of PLL6 and mutated each of its seven other pore-facing residues to Asn (Fig. 4C, D). Mutations at four of these tunnel facing residues (I703N, A705N, A706N, P717N) were able to restore LSB resistance to levels comparable to WT LetB or greater, and one additional mutant (G707N) restored partial resistance. Mutation of the final two positions (V708N, P701N) resulted in non-functional LetB variants. Thus, while the hydrophobic motifs appear to be critical (20/21 mutations are non-functional), other pore-facing hydrophobic residues are more tolerant of mutation (only 2/7 other mutations in PLL6 are non-functional).

Intriguingly, single amino acid substitutions at three of the above positions conferred even greater LSB resistance than WT LetB (I703N, A706N, P717N), and this level of resistance was comparable to that observed in the ΔRing6 mutant. The I703 side chain is tightly packed near the center of the pore-facing surface of PLL6 in the closed state of WT LetB, making van der Waals contacts with several other surrounding hydrophobic residues (V708, L711, and I714 of the same protomer, and V708 and L711 of the adjacent protomer; PDB: 6V0C) (Fig. S6A). Mutation of this residue would likely disrupt the tight hydrophobic packing, and may favor the open state where I703 is more loosely packed. A706 lies at the top of the hairpin portion of PLL6, and makes the only contacts between PLL6 and the neighboring Ring7 in the closed state (I834, in the short helical region of PLL7) (Fig. S6A), potentially helping to stabilize the conformation of PLL6. P717 lies at the end of the short helix of PLL6 and may be involved in helix breaking and creating a rigid connection to the core MCE domain. Ring6 and PLL6 have been shown to undergo the largest conformational changes in LetB, and may have a role in gating the tunnel via modulation of the Ring6 pore diameter. Thus, the increased LSB resistance of the ΔRing6 mutant may arise from the removal of this bottleneck in Ring6, and other mutations affecting Ring6/PLL6 may also increase LSB resistance by disfavoring the closed conformation of the tunnel.

The length and sequence of the PLLs varies considerably between MCE rings (Fig. 5F). To further explore how differences in sequence and length of the PLL at Ring6 may alter LSB resistance, we generated a series of mutants in which we replaced PLL6 with a PLL from a different MCE ring (Fig. 5A). We found that most of these mutants were indistinguishable from a strain completely lacking LetB in our phenotypic assay (Fig. 5B). But remarkably, replacement of PLL6 with PLL3 (6PLL3) greatly increased LSB resistance, to a degree greater than WT LetB and similar to the level that we observed for the ΔRing6 mutant (Fig. 5B). Like the ΔRing6 mutant, this 6PLL3 mutant conferred greater resistance to cholate than WT LetB (Fig. 5C). However, a 3PLL6 mutant (in which PLL3 is replaced with PLL6) did not fully relieve the exacerbated phenotype of a *pqiAB letAB* double mutant in the presence of LSB (Fig. 5D). All these mutants appeared to have a similar expression level in the complementation strains as detected by western blot (Fig. S6B). Using negative stain EM, we found that all of these LetB PLL6 replacement mutants assemble into the expected overall architecture (Fig. S6C). As most PLL swap mutants do not complement, yet properly fold, this suggests that swapping the PLLs disrupts the proper function of LetB. PLL2, PLL4, PLL5, and PLL7 are longer than PLL6 (≈24 residues as opposed to ≈17 residues), and modeling these longer PLLs into Ring6 suggests that this would lead to a steric clash between the tip of the loop and the C-terminal region of PLL7 in the next MCE ring (Fig. 5E). However, since these proteins still assemble into hexameric LetB tubes, accommodating these longer loops may restrict or even block the transport of substrates through the LetB tunnel at this location. In contrast, PLL3 is approximately the same length as PLL6 and would likely fill roughly the same volume in the LetB interior; yet because it has been grafted into a non-native location in Ring6 with low sequence identity to Ring3 (Fig. 5F), the contacts between PLL3 and surrounding residues in Rings5/6/7 are likely weaker and may make PLL3 in this context more flexible, and less able to adopt a rigidly closed state of the tunnel.

**Figure 5.**
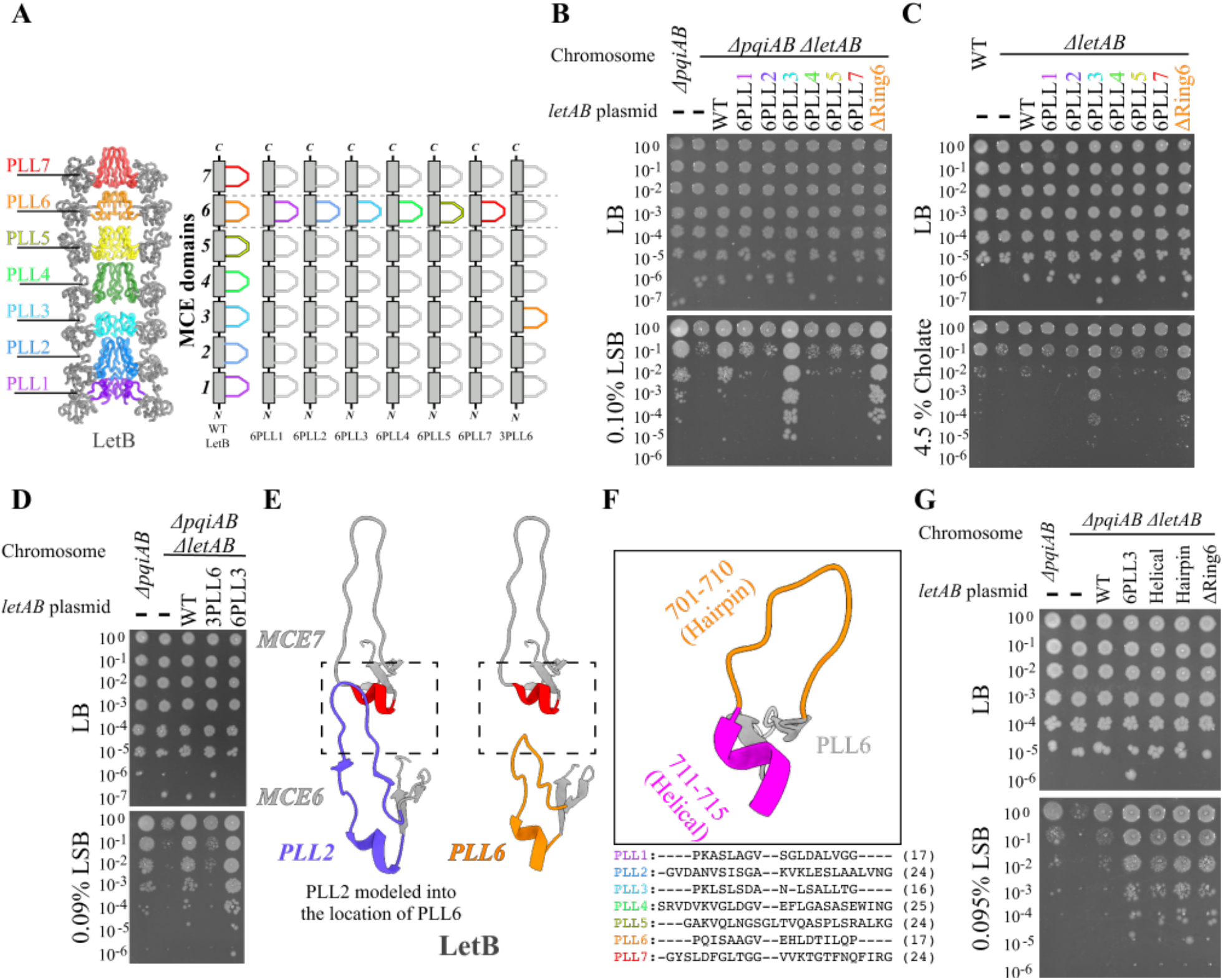
Phenotypes of mutants in which PLL6 is replaced with other PLLs. **(A)** Schematic of PLL6 swap mutants. PLLs 1-7 are shown as different colors, from purple to red. PLL6 was replaced with a PLL from each of the other MCE domains. **(B-D)** Cellular assay for the function of LetB mutants based on LSB resistance **(B and D)** or cholate resistance **(C)**. 10-fold serial dilutions of the specified cultures were spotted on LB agar containing any additives as indicated, and incubated overnight. **(E)** Illustration of the potential steric clash that would be induced if a longer PLL were introduced at MCE6. Modeling PLL2 into MCE6 leads to a longer PLL that would potentially clash with MCE7, while the tip of the original PLL6 is shorter and does not clash with MCE7. **(F)** PLL6 showing the short helical (magenta) and hairpin (orange) regions and sequence alignment of each LetB PLL. **(G)** Cellular assay for the function of LetB PLL6/PLL3 loop chimeras based on LSB resistance. 10-fold serial dilutions of the specified cultures were spotted on LB agar containing any additives as indicated, and incubated overnight.

To narrow down the region of PLL6 that modulates increased resistance to LSB and cholate compared to WT LetB, we replaced either the short helical segment or the hairpin segment of PLL6 with the corresponding region from PLL3 (Fig. 5F). Replacing only the hairpin led to increased LSB resistance comparable to the ΔRing6 mutant (Fig. 5G). Intriguingly, replacing only the short helical segment also led to increased LSB resistance. Thus, both the hairpin and helical region can independently modulate LSB resistance, suggesting that the properties of the PLL as a whole determine the ability of a LetB variant to confer LSB resistance, rather than a specific, localized site. Both of these PLL chimeras are expected to disrupt native interactions between PLL6 and the surrounding regions of the LetB tunnel, perhaps leading to increased flexibility and a reduced propensity to adopt the closed state.

## Discussion

Our combined results dissect the functional contribution of different MCE domains, rings and loops in the LetB protein and show how multi-domain MCE proteins like LetB can be engineered to remove an entire ring while preserving the overall structure. The ΔRing6 protein is not only highly stable and structurally similar to the WT, but surprisingly confers a higher level of LSB resistance. It is unclear if the higher resistance to LSB and cholate represents a hyper-active protein or a gain-of-function phenotype. As Ring6 is one of the most dynamic parts of LetB (Isom et al., 2020), and forms the narrowest region of the tunnel in the closed conformation, removing Ring6 may interfere with the ability of LetB to gate the transport tunnel and therefore result in a protein that is constitutively active. If Ring6 indeed serves as a gatekeeper to restrict transport through the tunnel, then other mutations that destabilize the closed state of the tunnel might be expected to also interfere with gating and result in increased LSB resistance. Indeed, while point mutations in conserved hydrophobic residues throughout the tunnel interior abolished LetB function, other mutations throughout PLL6 resulted in increased LSB resistance, comparable to removing Ring6 entirely. While the mechanism by which the mutations increase LSB resistance is not fully understood, all of them have the potential to increase the flexibility of PLL6 and/or disrupt interactions in the closed state, and thereby may shift the conformational equilibrium of LetB towards the open state, increasing flux of substrates through the tunnel. Similar conformational changes to those in Ring6 have also been observed in MCE rings of the orthologous protein, PqiB (Isom et al., 2020), suggesting that gating the tunnel by the PLLs may be a conserved mechanism by which multi-ring MCE proteins restrict transport.

## Materials and Methods

### Bacterial strains and plasmids

*E. coli* strains bBEL385, bBEL384, and bBEL466, with deletions of *pqiAB* (*ΔpqiAB*), *pqiAB* and *letAB* (*ΔpqiAB ΔletAB*), or *letAB* (*ΔletAB*) respectively, are derivatives of BW25113 and were reported previously (Isom et al., 2017). Strains were subjected to whole genome sequencing to confirm their genotype. Plasmid pBEL1620 (pET17b-*yebST*, Addgene #175804 (Isom et al., 2020, 2017) encoding the complete WT *letAB* operon was used to complement the *ΔpqiAB ΔletAB* strain. Plasmids were transformed into bBEL384 or bBEL466 using the CaCl_2_ heat shock method. Bacteria were routinely grown at 37°C in LB media (Difco) or on LB Agar plates for plasmid propagation. For all strains harboring a plasmid, the liquid medium and/or plate was supplemented with 100 μg/ml carbenicillin, unless otherwise noted.

To test the impact of mutations in LetB, the *letB* region of pBEL1620 (pET17b-*yebST*) was mutated using Gibson assembly (Gibson et al., 2009). For MCE ring deletions, we used domain boundaries similar to those previously described for engineering shorter LetB proteins (Isom et al., 2020), which include the core MCE domain and C-terminal linker (see Table S2). For PLL grafting experiments, we defined the boundaries of each PLL based upon the location of conserved proline and glycine residues near each end (e.g., Pro346 and Gly361 in PLL3) and swapped out the remaining residues which defined the PLL for that ring in order to assay PLL function. To probe the importance of the conserved hydrophobic motif ΦxxΦΦ in each PLL, we mutated each residue to Asn. Leu is the most common residue at these sites, mutation to Asn approximately preserves residue size and shape, but converts the hydrophobic residue to a polar one.

### Bacterial growth assays

LSB (Sigma-Aldrich, catalog #D0431) and cholate (Alfa Aesar, catalog #A17074-18) sensitivity of *E. coli* strains was based upon a previously described growth assay on LB agar plates (Isom et al., 2020, 2017). To improve discrimination between WT and mutant strains, we systematically explored two variables that we suspected may modulate the phenotype: 1) the preparation of the LB agar; and 2) the growth phase of the cultures used for dilutions and plating.

The manner in which the LB Agar was prepared had a significant effect on level of LSB and cholate sensitivity of *E. coli* strains, and consequently on the optimal concentrations of LSB and cholate required; similar observations have been previously reported for other bacterial growth phenotypes (Kolich et al., 2020). We prepared LB agar from separate components (“LB Agar 1”: 10g Tryptone (Gibco, catalog #211705, 10g NaCl (Sigma-Aldrich, catalog #S3014), 5g Yeast extract (Gibco, catalog #212750, 15g Agar (BD Difco, catalog #214530) per 1L H_2_O); by combining pre-mixed LB broth (BD Difco, catalog #244620) with agar (“LB Agar 2”); or a complete LB+agar pre-mix (“LB Agar 3”; BD Difco, catalog # DF0445-07-6) and found that LB Agar 3 requires about ≈10x less of the additive lauryl LSB (≈0.1%) or ≈2x less cholate (≈4.5-5%) than LB Agar 1 to achieve the same degree of bacterial growth inhibition (Fig. S1F). Unless otherwise noted, all genetic complementation assays reported in this work were conducted using LB Agar 3. We also tested the impact of the growth phase of the starting culture on the relative sensitivity of WT and mutant strains (Fig. S1G) and unless otherwise noted all genetic complementation assays reported in this work were conducted as described below. Stock solutions of LSB in water (5% w/v), cholate (40% w/v), and bile salts (40% w/v; Sigma-Aldrich, catalog #B8756) were prepared and stored at -80°C. Stock solutions of vancomycin (50 mg/ml; Sigma-Aldrich, catalog #V2002) and rifampicin (100 μg/ml; MedChemExpress, catalog #HY-B0272) were stored at -20°C. As the sensitivity phenotypes were sensitive to relatively small changes in concentration, a range of concentrations were used for each experiment from 0.09% to 0.11% LSB, 4.5% to 6% cholate, 0.5% to 1.5% bile salts, 180 μg/ml to 200 μg/ml vancomycin, and 2 μg/ml to 5 μg/ml rifampicin.

Overnight cultures grown in LB were diluted 1:50 into fresh LB without antibiotics. Cultures were grown for ≈1.5 hours at 200 rpm and 37 °C until reaching an OD600 of ≈1.0, then normalized to a final OD600 of 1.0 with fresh LB. From these normalized cultures, 10-fold serial dilutions in LB were prepared in a microtiter plate, and 1.5ul of each dilution was spotted onto plates containing LB Agar 3 alone, or LB Agar 3 supplemented with LSB or cholate. Plates were incubated ≈20-24 hours at 37°C and then photographed using a ChemiDoc XRS+ System (Bio-Rad). At least three independent transformants were used to perform replicates for phenotypic assays.

### Live-cell Imaging

As the *ΔpqiAB ΔletAB* growth defect in the presence of LSB is highly sensitive to LB/Agar formulations, we first validated that the growth defect of a *ΔpqiAB ΔletAB* strain in the presence of LSB was reproducible on LB/Agarose, which is used in all our microscopy experiments (Fig. S1F), and found that LSB concentrations between 0.1% and 0.5% LSB substantially inhibited *E. coli* growth.

Overnight cultures grown in LB at 37°C were diluted 100x and grown at 37°C for ≈60 minutes at 200 rpm until they reached an OD600 of ≈0.6-0.8 for logarithmic phase. Propidium Iodide dissolved in water (Sigma-Aldrich, catalog #P4864-10ML) was then added to *E. coli* cells at a final concentration of 5 μg/ml. 1 µl of cells/PI mix were pipetted onto agar pads composed of 1.5% LB Agarose, supplemented with either 0.2% LSB, 5% cholate, or 38 μg/ml of chloramphenicol for additive conditions, on glass depression slides (United Scientific Supplies, catalog #470235-728) and sealed with a #1.5 18 × 18 mm coverslip (Chemglass Life Sciences, catalog # CLS-1764-1818) and petroleum jelly (Vaseline). The time between the initial exposure of cells to agarose pads to the time at which image acquisition was started was recorded and took on average ≈3-5 minutes. Live cells were imaged for approximately 4 hours in an environmental chamber which was set at 37°C. Images were collected on a Nikon Eclipse Ti microscope with a Nikon 60x N.A. 1.4 oil immersion Plan Apochromat Ph3 phase-contrast objective lens with an Andor Zyla 5.5 megapixel sCMOS camera. Exposures were acquired every minute for 240 minutes with 3×3 binning for phase contrast imaging. PI imaging was performed with epifluorescence microscopy with an electron multiplying charged-coupled device camera (EMCCD) at an emission wavelength of 640 nm and exposure time of 200ms at ≈6% laser power with 3×3 binning. Three independent cultures of *E. coli* were used to perform three replicates for each condition. For each imaging condition between 15 to 20 independent fields of view (FOV)s were captured to visualize approximately the same number of cells for each replicate.

Time-lapse movies were loaded into Fiji (Schindelin et al., 2012) or ImageJ (Schneider et al., 2012) for subsequent analyses. Images were manually thresholded in Fiji and the percent of PI+ cells in each FOV were counted.

### Expression and purification of membrane associated LetB

Starting with plasmid pBEL1284, which encodes an N-terminal 6xHis2xQH-TEV tagged LetA and untagged LetB, mutant forms of LetB were created by introducing the target mutations into pBEL1284 by oligo-based mutagenesis and subsequent Gibson assembly of the resulting PCR products. The resulting expression plasmids were transformed into T7 Express *E. coli* cells (NEB).

For protein production, overnight cultures of the resulting strains were grown in LB supplemented with 1% glucose, diluted 1:50 into fresh LB and grown at 37°C with shaking at 200 rpm to an OD600 of ≈0.8-1.0, then induced with 0.2% arabinose for 4 hours at 37°C. Cultures were harvested by centrifugation and cells were then resuspended and lysed in Buffer A (50 mM Tris pH 8.0, 300 mM NaCl, 10% glycerol) by two passes through an Emulsiflex-C3 cell disruptor (Avestin). Lysates were clarified at 15,000g for 30 minutes and membrane fractions were isolated by ultracentrifugation at 180,000g for 45 minutes. Membranes were solubilized for one hour in 50 mM Tris pH 8.0, 300 mM NaCl, 10% glycerol, 25 mM DDM by rocking at 4°C, after which insoluble debris was pelleted by ultracentrifugation at 180,000g for 45 minutes. Solubilized membranes were then incubated with Ni beads (Ni Sepharose Fast Flow, GE Life Sciences) for 30 minutes at 4°C and purified by immobilized metal affinity chromatography (IMAC) with intermediate washes of Buffer A (50 mM Tris pH 8.0, 300 mM NaCl, 10% glycerol, 0.5 mM DDM) supplemented with 20 mM imidazole for the first wash and 40 mM imidazole for the second wash, before eluting from the Ni beads with Buffer A supplemented with 250 mM imidazole pH 8.0. LetAB mutants were further purified by size exclusion chromatography (Superdex 200 Increase 10/300 GL column) on an ÄKTA pure system in 20 mM Tris pH 8.0, 150 mM NaCl, 0.5 mM DDM, and 10% glycerol. Gel filtration fractions were then added directly to an EM grid (see Electron microscopy grid preparation section of methods).

### Expression and purification of ΔRing6 LetB for cryoEM

For cryo-EM, the ΔRing6 LetB periplasmic domain (pBEL2004) was transformed into Rosetta (DE3) cells (Novagen) for protein expression and overnight cultures were grown in starter cultures supplemented with 1% glucose overnight at 37°C. Overnight cultures were subsequently diluted 100x into LB and 1000x metals and upon reaching an OD600 of ≈0.8 cells were induced with 1 mM IPTG for 4 hours at 37°C. Cultures were harvested by centrifugation and cells were resuspended and lysed in 20 mM Tris pH 8.0, 300 mM NaCl by two passes through an Emulsiflex-C3 cell disruptor (Avestin) and lysates were clarified by centrifuging at 30,000g for 30 minutes and the supernatant was incubated with Ni resin (Ni Sepharose Fast Flow, GE Healthcare) for 30 minutes at 4°C. ΔRing6 was purified by immobilized metal affinity chromatography (IMAC) with 20 mM Tris pH 8.0, 150 mM NaCl, 250 mM imidazole pH 8.0 as elution buffer. Elution fractions were further purified with size exclusion chromatography (Superdex 200 Increase 10/300 GL column) on an ÄKTA pure system in 20 mM Tris pH 8.0, 150 mM NaCl. Gel filtration fractions containing purified ΔRing6 LetB were concentrated to ≈1mg/ml for electron microscopy grid preparation.

### Western blotting

*LetB* expression levels were assessed using previously established methods (Isom et al., 2020, 2017). 10 mL cultures of *E. coli* strains Δ*pqiAB*, Δ*pqiAB* Δ*letAB*, and Δ*pqiAB* Δ*letAB* containing each complementation plasmid was grown to an OD of 0.6-0.8. The cells were pelleted at 4000 g for 10 mins and resuspended in 1 mL of freeze thaw lysis buffer (PBS, 1 mg/ml lysozyme, 0.5 mM EDTA and 1 μl/ml of benzonase (Millipore, catalog #70746-3)), and incubated on ice for 1 hour. The samples underwent eight cycles of freeze-thaw lysis by alternating between liquid nitrogen and a 37°C heat block. After lysis, the cells were centrifuged at 23,000 g for 15 minutes in a microcentrifuge and the membrane-containing pellets were resuspended in 40 μl of 2xSDS-page loading buffer. 10 μl samples were separated on an SDS-PAGE gel and transferred to a nitrocellulose membrane. The membranes were blocked in PBST containing 5% powdered milk for one hour. The membranes were then incubated with primary antibody in PBST + 5% powdered milk (either rabbit polyclonal anti-LetB rabbit (dilution of 1:10,000) or polyclonal anti-BamA (dilution of 1:2000)) overnight at 4°C (Isom et al., 2017). The membranes were then washed 3 times with PBST and were incubated for 1 hour with 1:10,000 IRDye 800CW Goat anti-rabbit IgG Secondary Antibody (LI-COR, Catalog # 926-32211) in PBST + 5% BSA. Membranes were then washed 3 times with PBST and imaged on a LI-COR Odyssey Classic.

### Electron Microscopy grid preparation and data collection

Negative stain grids were prepared by diluting protein samples onto Formvar/Carbon 400 mesh on Copper grids (Ted Pella Inc.) which were previously glow discharged in a Pelco easiGlow Discharge Unit (Ted Pella Inc.) with a 30 second hold time and 30 second glow discharge time. 3 µl of protein sample diluted to a concentration of between 0.02 mg/ml and 0.05 mg/ml was added to a grid and then manually blotted away with Whatman Grade 1 filter Paper (GE Healthcare). 3 µl of 0.75% Uranyl Formate was then added to the grid and blotted away, 5 times. Grids were then imaged in a Talos L120C TEM (FEI) at 73,000 magnification with a pixel size of 2.00 Å/pixel.

ΔRing6 LetB cryo-EM samples were prepared by concentrating ΔRing6 LetB after size exclusion to 0.75 mg/ml. 300 Cu mesh quantifoil 1.2/1.3 grids were used and grids were glow discharged in a Pelco easiGlow Discharge Unit (Ted Pella Inc.) with a 30 second hold time and 12 second glow discharge time. Grids were then prepared using a FEI Vitrobot Mark IV with 3 second blot time and blot force of 5 and screened on the NYU Talos Arctica using Leginon (Carragher et al., 2000). Complete datasets were collected at the Pacific Northwest Center for Cryo-EM (PNCC) on a Titan Krios using SerialEM (Mastronarde, 2003) equipped with a K3 camera and set to a pixel size of 0.636Å (super resolution pixel size: 0.318Å). Data collection parameters can be found in Table S1.

### Electron microscopy data processing

The data processing workflow is shown in Fig. S4. A combination of RELION 3.0, RELION 3.1, and cryoSPARC were used. Motion correction was carried out using RELION’s implementation of MotionCor2 (Zheng et al., 2017) and CTF estimation was performed using CTFFIND4 (Rohou and Grigorieff, 2015). Particle picking was done using cryoSPARC’s blob picker function (Punjani et al., 2017) followed by cryoSPARC’s template based picker. 2D classification was carried out to assess particle quality and curate a final stack of 608,911 particles. An *ab initio* model was generated with the full particle set in cryoSPARC. Homogenous refinement was carried out in cryoSPARC. Particles corresponding to the best map were imported into RELION 3.1 (Zivanov et al., 2018) where CTF and aberration refinements (Zivanov et al., 2020) were done, followed by 3D refinement. Iterative 3D classification was performed in RELION as shown in Fig. S4, resulting in the highest resolution map for the full protein complex (Map 1, EMDB EMD-25066). In this map, Ring 1 is in the closed conformation. Additional 3D classification on the full map did not yield additional conformations or higher resolution.

In order to improve the local resolution and analyze conformational heterogeneity, signal subtraction, 3D classification and refinement were carried out in RELION. Masks of three consecutive MCE rings were used for signal subtraction. Other masking and signal subtraction strategies were attempted, but masking three consecutive rings led to the highest resolution maps. One map was resolved (Map 2, EMDB EMD-25067) in which a conformational change was observed in Ring1, leading to the open state of Ring1 (similar to PDB 6V0J), Other maps did not have any apparently different features from Map 1 or Map 2. We varied the number of classes and regularisation parameter T (Scheres, 2012) and found that classifying into a larger number of classes (12) and high T values (T=100) resulted in classification of Ring 1into the open and closed states.

The resolution of the best maps varied from ≈3.2 to ≈3.6Å and for each map the overall resolution was estimated using the gold-standard Fourier Shell Correlation criterion (FSC = 0.143). The 3D FSC for the reported maps (Fig. S5B) were generated at https://3dfsc.salk.edu/ (Tan et al., 2017). Refinements were carried out with C6 symmetry imposed or no symmetry imposed (C1) for all maps used in model building. Local resolution maps were computed in RELION (Fig. S5A). Final maps are with C6 symmetry imposed. Each map was sharpened using global B-factor sharpening of -98 Å^2^ in RELION (Zivanov et al., 2018) and density modification was performed using the resolve_cryo_em algorithm implemented in phenix (Terwilliger et al., 2020) or DeepEMhancer (Sanchez-Garcia et al., 2020). These modified maps together with the unmodified map directly from refinement were used in the model building process.

### Model Building

The periplasmic domain WT LetB (PDB ID: 6V0D) (Isom et al., 2020) was used as a starting model. MCE domain 6 was deleted from the LetB model and MCE domain 7 was treated as a rigid body and fit into corresponding density in the ΔRing6 map. For map number 2, corresponding to the open state of Ring1, the corresponding open state of Ring 1 from WT LetB was used as a starting model (PDB ID: 6V0J) (Isom et al., 2020). Residues were manually adjusted using COOT (Emsley et al., 2010). The models were then subjected to the real_space_refine algorithm in PHENIX (Afonine et al., 2018). Iterative rounds of manual model building in coot and refinement in PHENIX were carried out. Three to five refinement cycles were used, optimizing the non-bonded weight parameter and adding riding hydrogens from phenix.reduce (Word et al., 1999) to optimize the fit and reduce clashes. Models corresponding to maps #1 (PDB ID: 7SEE; EMD-25066) and #2 (PDB ID: 7SEF; EMD-25067) were deposited in the PDB.

For validation, statistics regarding the final models (Table S1) were derived from real_space_refine algorithm of phenix and MolProbity (Williams et al., 2018), EMRINGER (Barad et al., 2015), and CaBLAM (Williams, 2015) from the PHENIX package (Adams et al., 2010) and were used for model validation. Model correlations to our EM maps were estimated with CC calculations from the PHENIX package (Adams et al., 2010), as well as the Q-score plugin for Chimera (Fig. S5C; (Pintilie et al., 2020)). CHAP (Klesse et al., 2019) was used to calculate the radii of the WT LetB and the ΔRing6 mutant, and the radius profile of the permeation pathway was visualized and plotted in MATLAB. Model visualization and analyses were performed with either the UCSF Chimera package (Pettersen et al., 2004), UCSF ChimeraX package (Goddard et al., 2018; Pettersen et al., 2021), or the Pymol Molecular Graphics System (version 2.4.0 Schröodinger, LLC). The PyMOL plugin RmsdByResidue (https://pymolwiki.org/index.php/RmsdByResidue) and visual inspection was used to assess the greatest differences between ΔRing6 and WT LetB (PDB: 6V0C & 6V0D).

## Supporting information

Associated Map/Model #1 & #2

## Accession numbers

The cryo-EM maps for the full length map with a closed state in Ring1 (map 1) and the open state of Ring1-3 (map 2) for the ΔRing6 mutant have been deposited in the Electron Microscopy Data Bank with accession codes EMDB: **<u>EMD-25066</u>** (map 1) and EMDB: **<u>EMD-25067</u>** (map 2). The coordinates of the atomic models have been deposited in the Protein Data Bank under accession codes PDB: **<u>7SEE</u>** (model 1) and PDB: **<u>7SEF</u>** (model 2). Cryo-EM data for the ΔRing6 mutant LetB were deposited in EMPIAR: **<u>47483752</u>**. Plasmids have been deposited in Addgene, and identifiers are in Table S2. Coordinates and maps are available for immediate download as supplementary files associated with this preprint, and will be publicly released from the PDB/EMDB as soon as the entries are processed.

## Acknowledgments

We would like to thank Rachel Redler, Pattana Jaroenlak, Joseph Sudar, Noelle Antao, and Mark MacRae for critical reading and feedback on our manuscript and the members of the Bhabha/Ekiert labs, NYU MSTP program, and NYU Department of Microbiology for helpful discussions; Janette Myers and the staff at PNCC, as well as Bill Rice and Bing Wang at NYU Langone Health’s Cryo–Electron Microscopy Laboratory (RRID: SCR_019202) for assistance in microscope operation and data collection; Ian Henderson (The University of Queensland), for providing antibodies against BamA and LetB; Kristen Dancel-Manning and Fengxia Lang for overseeing use of the TalosL120C electron microscope and Michael Cammer and Yan Deng for light microscopy training as part of NYU Langone’s Microscopy Laboratory (RRID: SCR_017934) supported by the Cancer Center Support Grant P30CA016087 at the Laura and Isaac Perlmutter Cancer Center; Central Lab Services team at the Skirball Institute supervised by Michael Mirabile for preparation of media and buffers in large quantities; and the HPC team for high-performance computing support. We gratefully acknowledge the following funding sources: R00GM112982 (NIH/NIGMS, to G.B.), DFS-20-16 (Damon Runyon Cancer Research Foundation, to G.B.), R35GM128777 (NIH/NIGMS, to D.C.E.), 1F30AI154907 (NIH/NIAID, to C.V.).

## Author Contributions

CV.: Conceptualization, data curation, investigation, writing - original draft, review and editing. N.C.: Validation, supervision, writing - review and editing. G.L.I.: Investigation, conceptualization, writing - review and editing. G.B.: Supervision, conceptualization, project administration, funding acquisition, writing - original draft, review and editing. D.C.E.: Supervision, conceptualization, project administration, funding acquisition, writing - original draft, review and editing.

---

**Figure S1.**
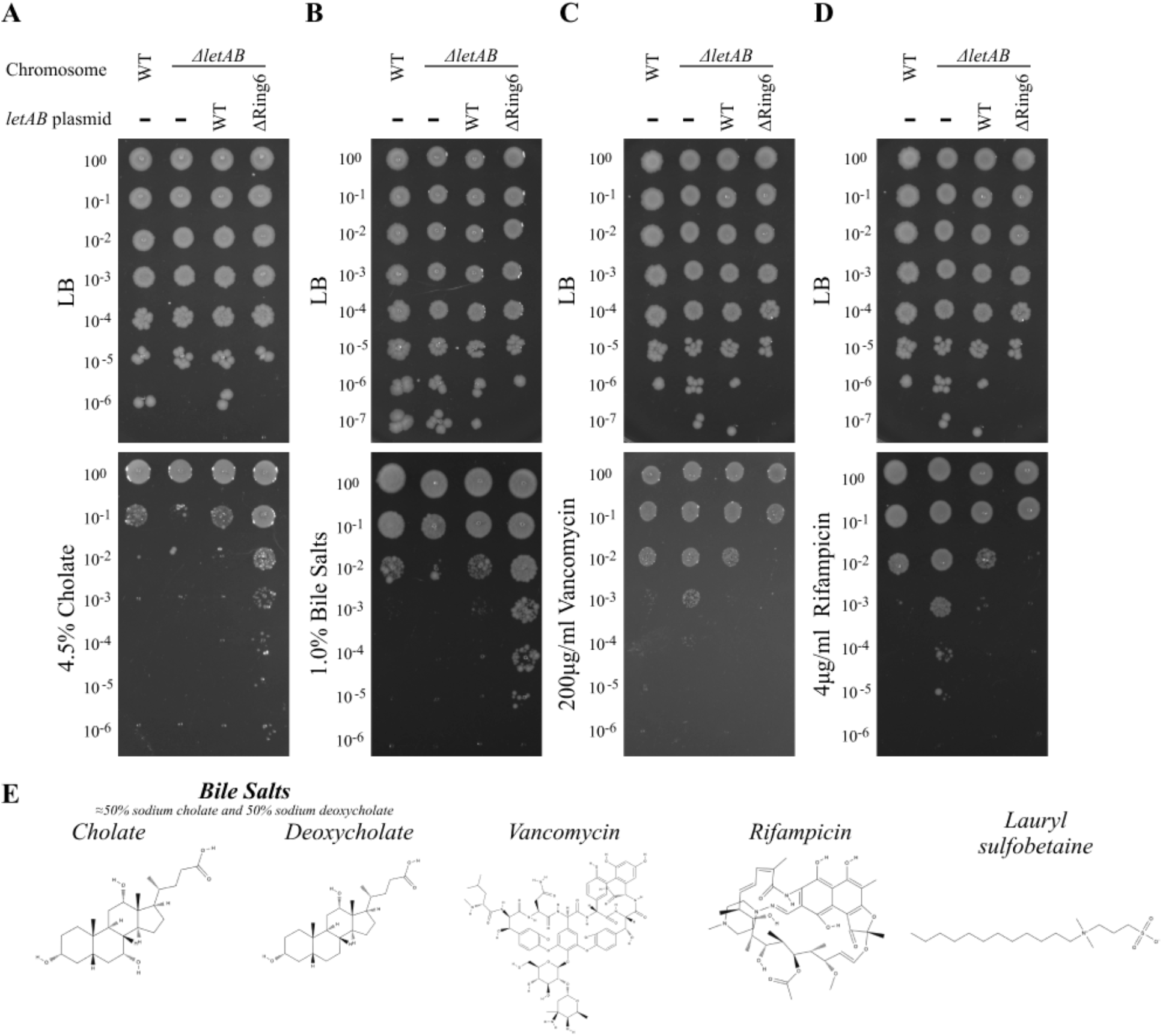
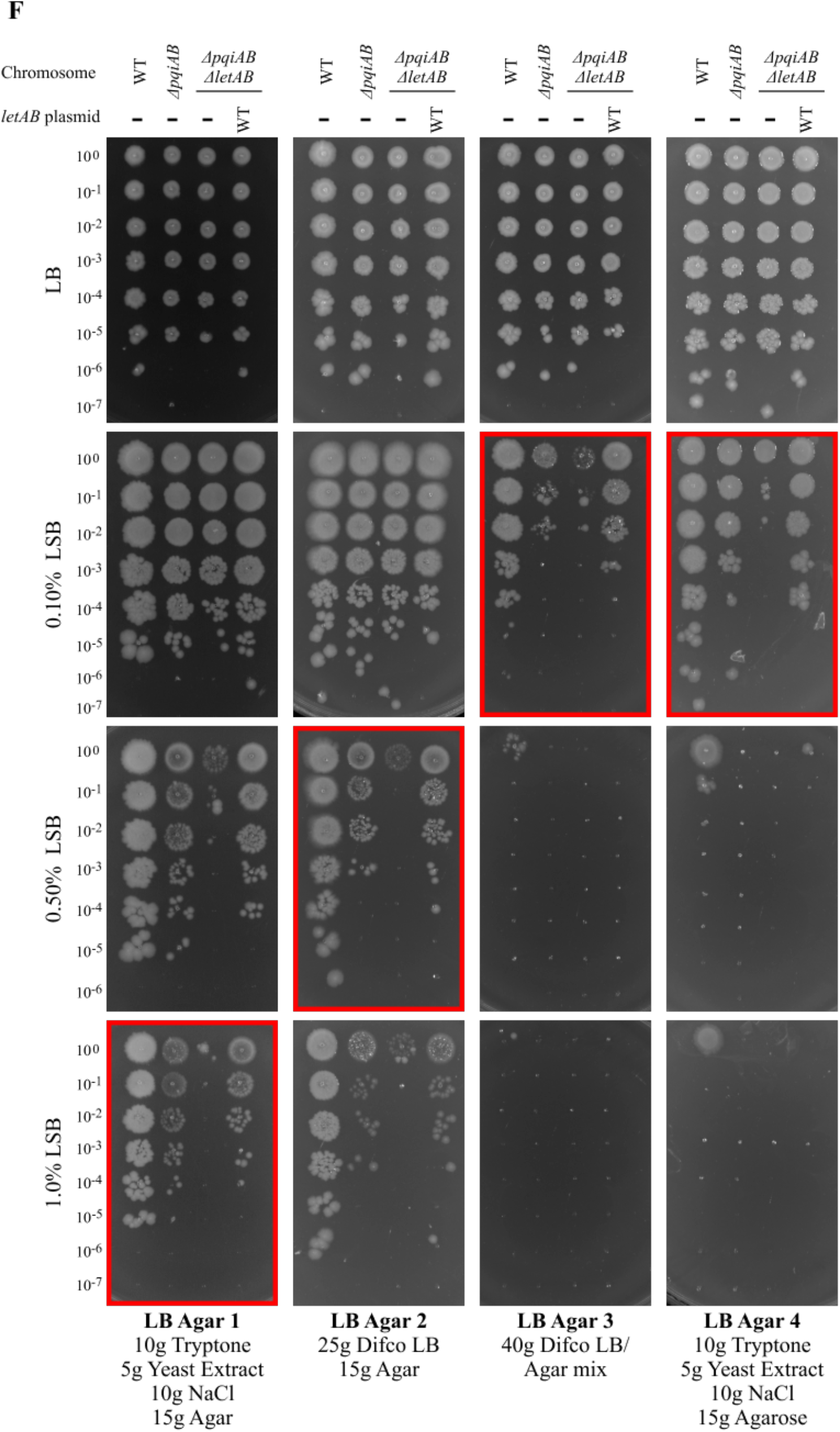
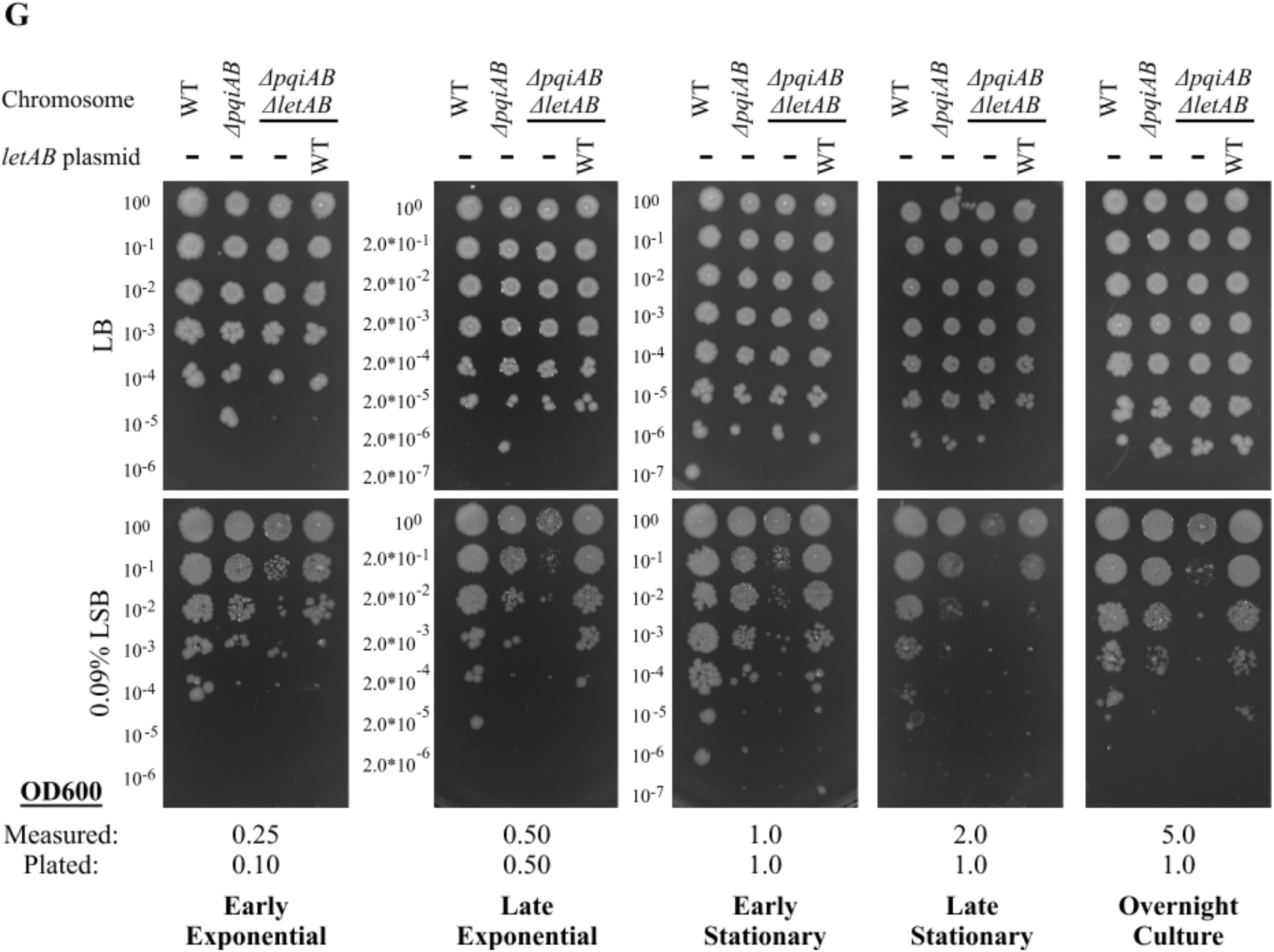
*LetB* phenotypes. The *ΔletAB* strain grows slightly less well than the WT strain in the presence of 4.5% cholate **(A)** and 1.0% bile salts **(B)**, and is slightly more resistant compared to the WT strain in the presence of 200 μg/ml vancomycin **(C)**, and 4 μg/ml rifampicin **(D)**. The *ΔletAB* strain can be complemented by the expression of WT *letAB* from a plasmid to restore WT growth levels. ΔRing6 mutant also shows increased resistance to cholate and bile salts, and decreased resistance to vancomycin and rifampicin. 10-fold serial dilutions of the specified cultures were spotted on LB agar containing any additives as indicated, and incubated overnight. **(E)** The chemical structure of the additive used in each phenotypic assay is shown. **(F)** Influence of agar/media formulation on LetB LSB resistance phenotype. 10-fold serial dilutions of the specified cultures were spotted on LB agar containing any additives as indicated, and incubated overnight. Optimal LSB conditions for each agar/media formulation are boxed in red. **(G)** Indicated strains plated at different growth phases and densities. All plates are labelled with the approximate OD600 of the four strains at the time they were plated, as well as the OD600 to which each strain was normalized to before plating. Serial dilutions of the indicated cultures were spotted on LB agar and incubated overnight.

**Figure S2.**
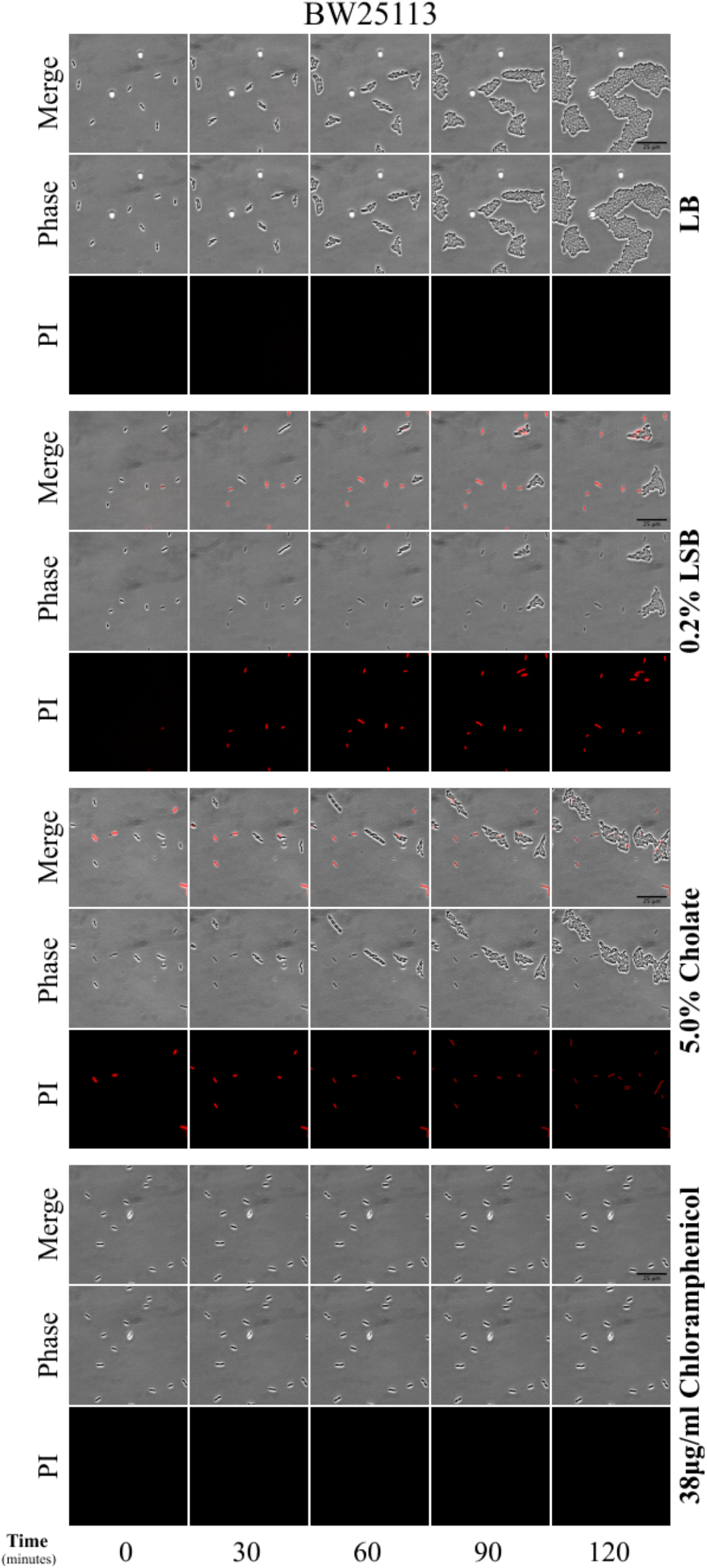
Unmerged images of phase contrast and propidium iodide channels. Images are the same as those shown in Fig. 1C.

**Figure S3.**
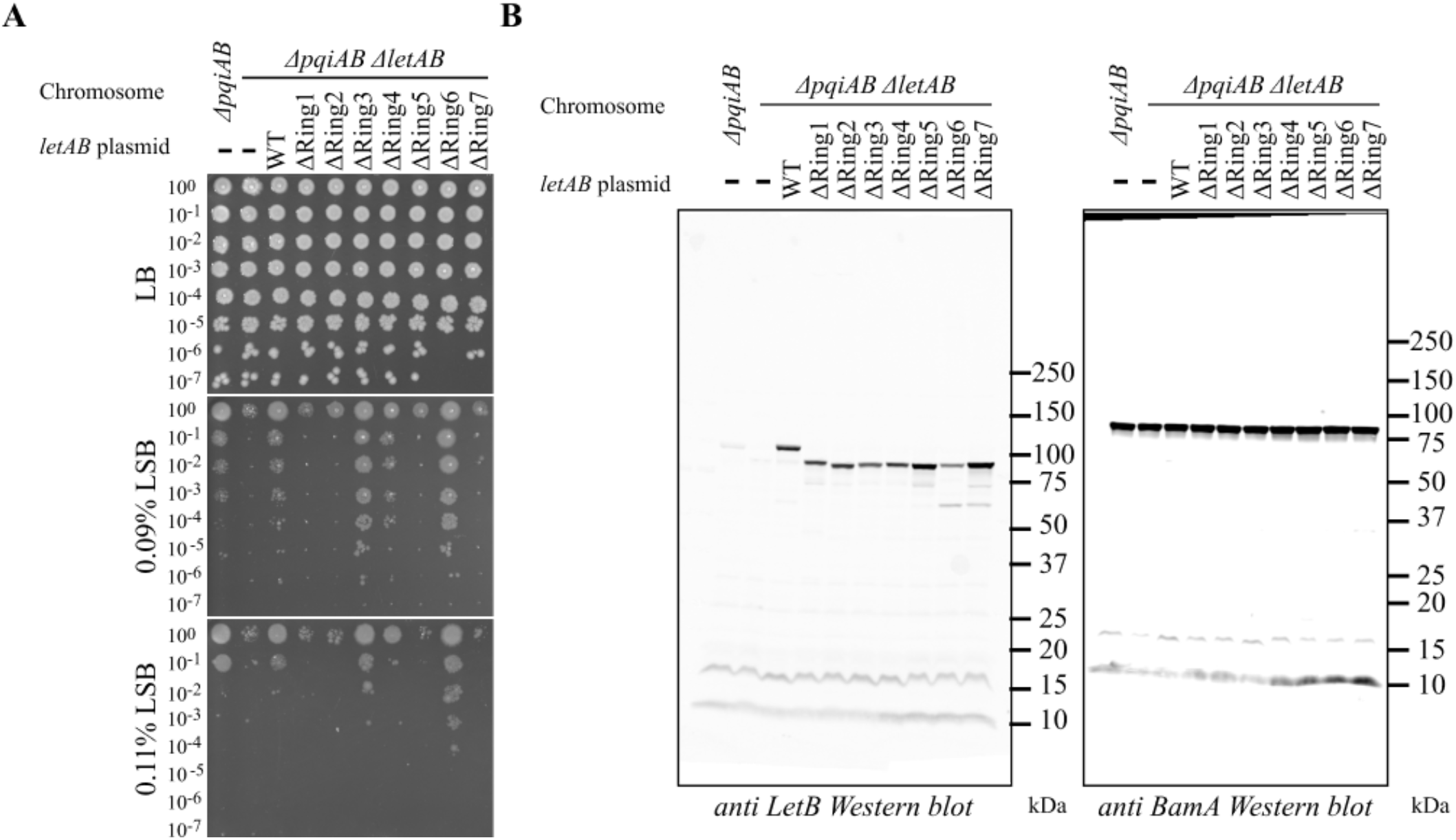
Additional data related to Fig. 2. **(A)** Cellular assay for the function of LetB ring deletion mutants based on LSB resistance. 10-fold serial dilutions of the specified cultures were spotted on LB agar containing any additives as indicated, and incubated overnight. Lower LSB concentration (0.09%) more clearly highlights the differences between the parental and knockout strains to differentiate strains that complement like WT, while a higher LSB concentration (0.11%) more clearly highlights the increased resistance of ΔRing6. The LB and lower LSB concentration are the same images as in Fig. 2B. **(B)** Uncropped lanes of western blot shown in Fig. 2D.

**Figure S4.**
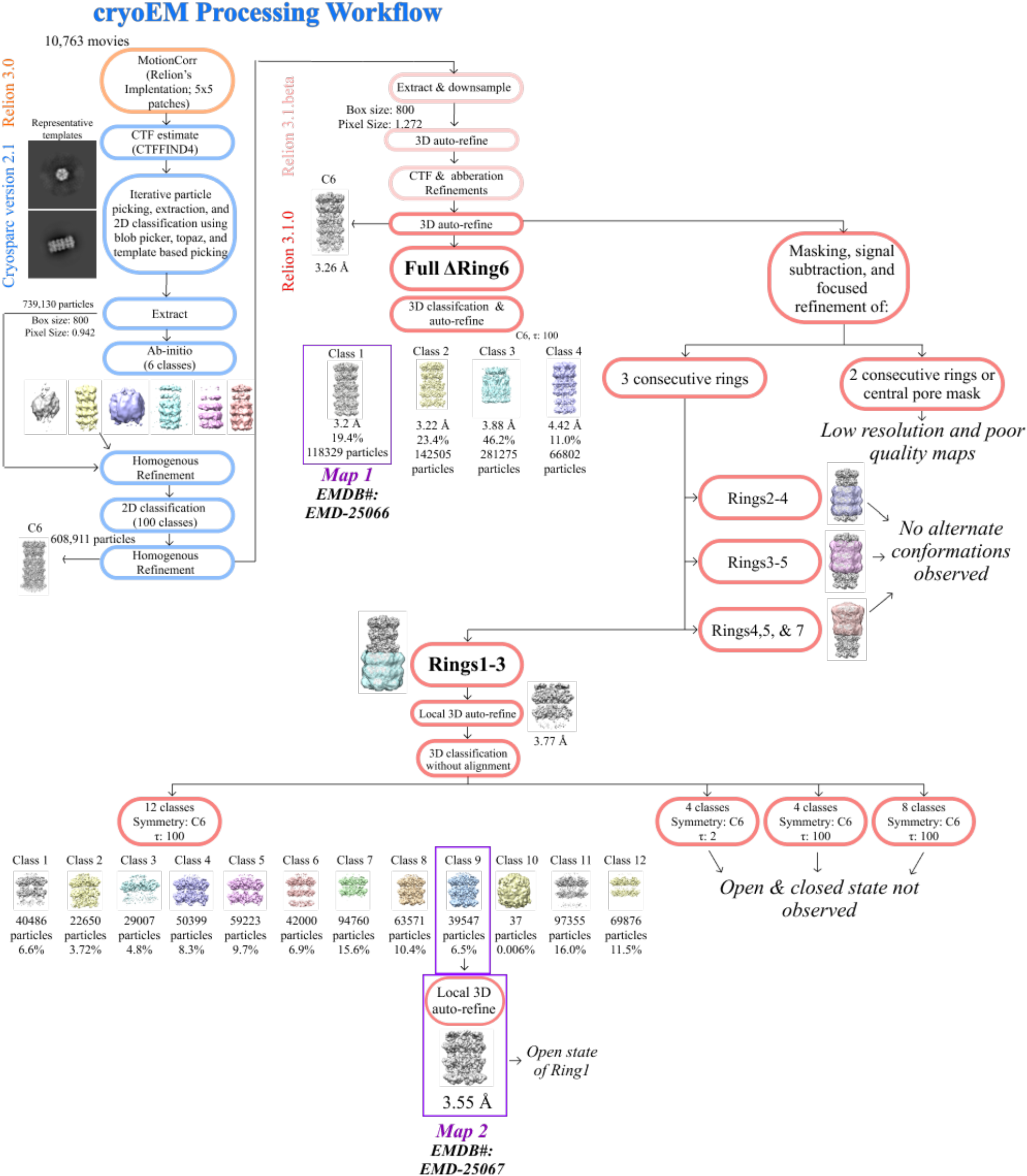
Cryo-EM data processing workflow for ΔRing6 LetB mutant.

**Figure S5.**
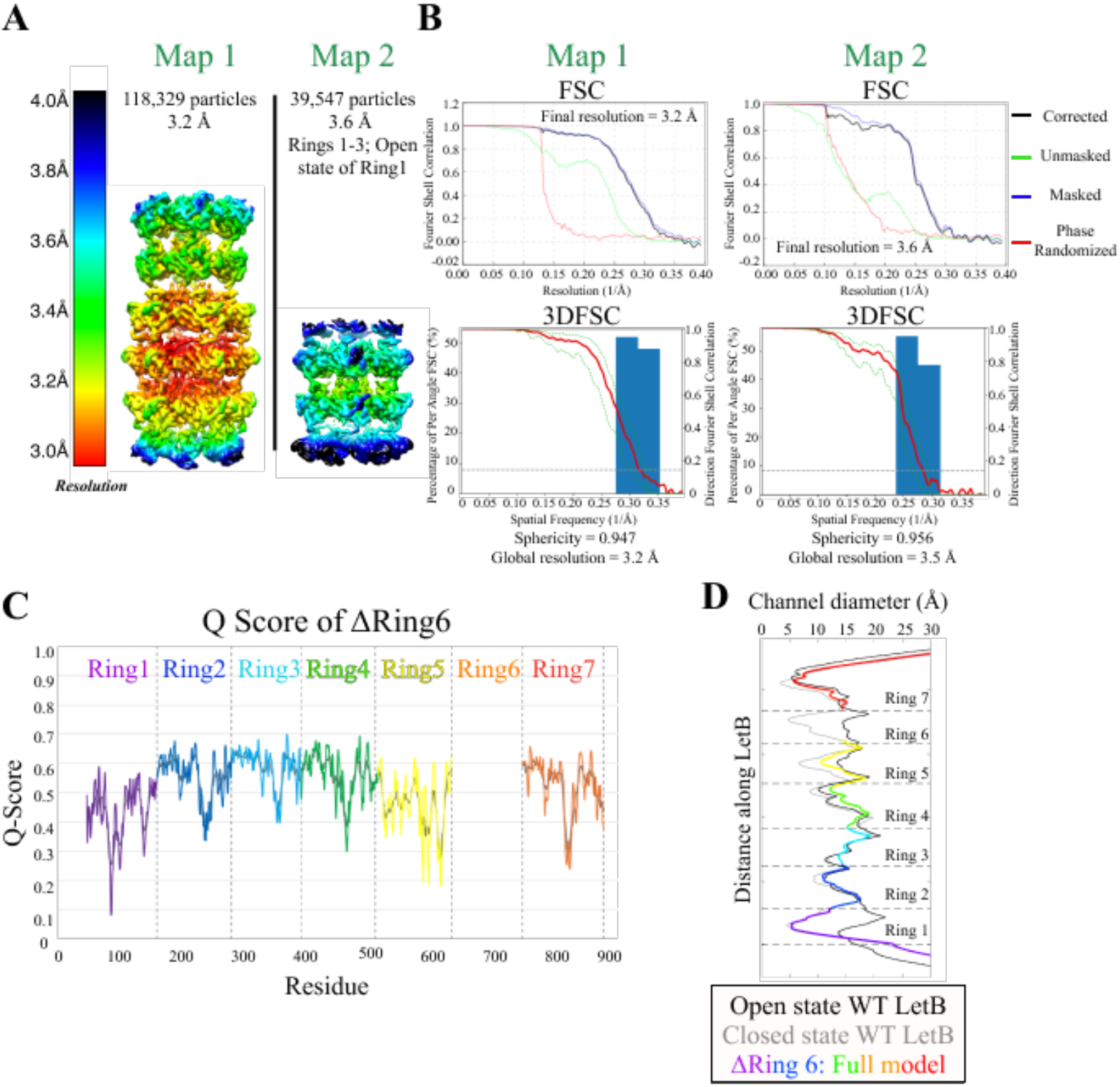
Additional data for cryo-EM structure. **(A)** Density maps of ΔRing6 LetB colored by local resolution, using local resolution estimation in RELION. **(B)** Fourier Shell Coefficient (FSC) from RELION and 3D FSC curves for Map1 and Map2, using the 3DFSC processing server (Tan et al., 2017). **(C)** Qscore validation plot from Chimera (Pintilie et al., 2020) with the Q score per residue plotted for the one residue average (black) and the five residue average (rainbow). **(D)** Diameter of the ΔRing6 mutant (model #1; rainbow) overlaid with the diameter of the WT LetB tunnel in the open (black) and closed (gray) states, measured using CHAP (Klesse et al., 2019).

**Figure S6.**
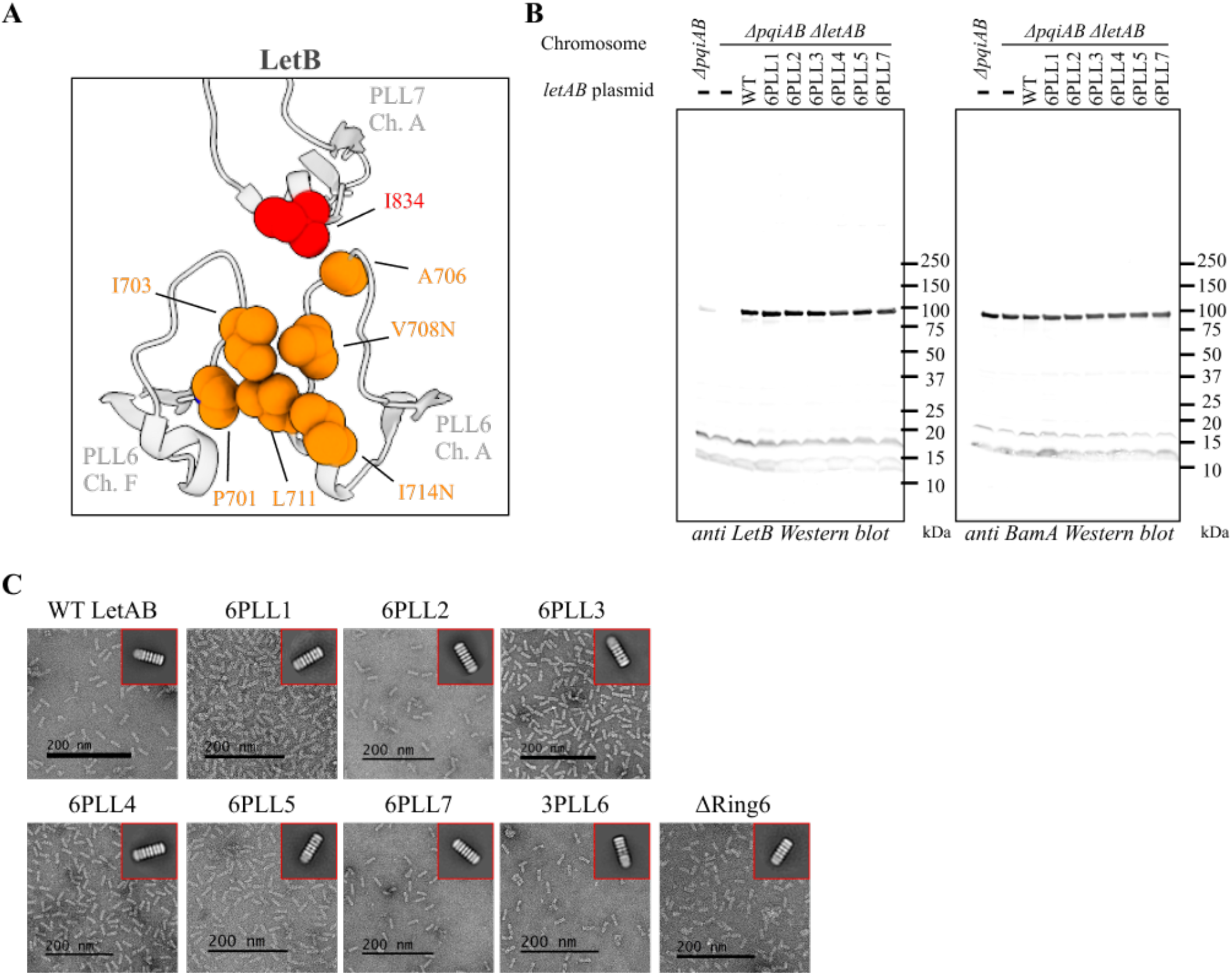
Additional characterization of PLL6 mutants. **(A)** PLL6 and the van der Waals contacts with surrounding residues (PDB ID: 6V0C). **(B)** Uncropped lanes of western blots shown in Fig. 5B. **(C)** Example negative stain micrographs and 2D classes of mutants as indicated.

**Table S1:**
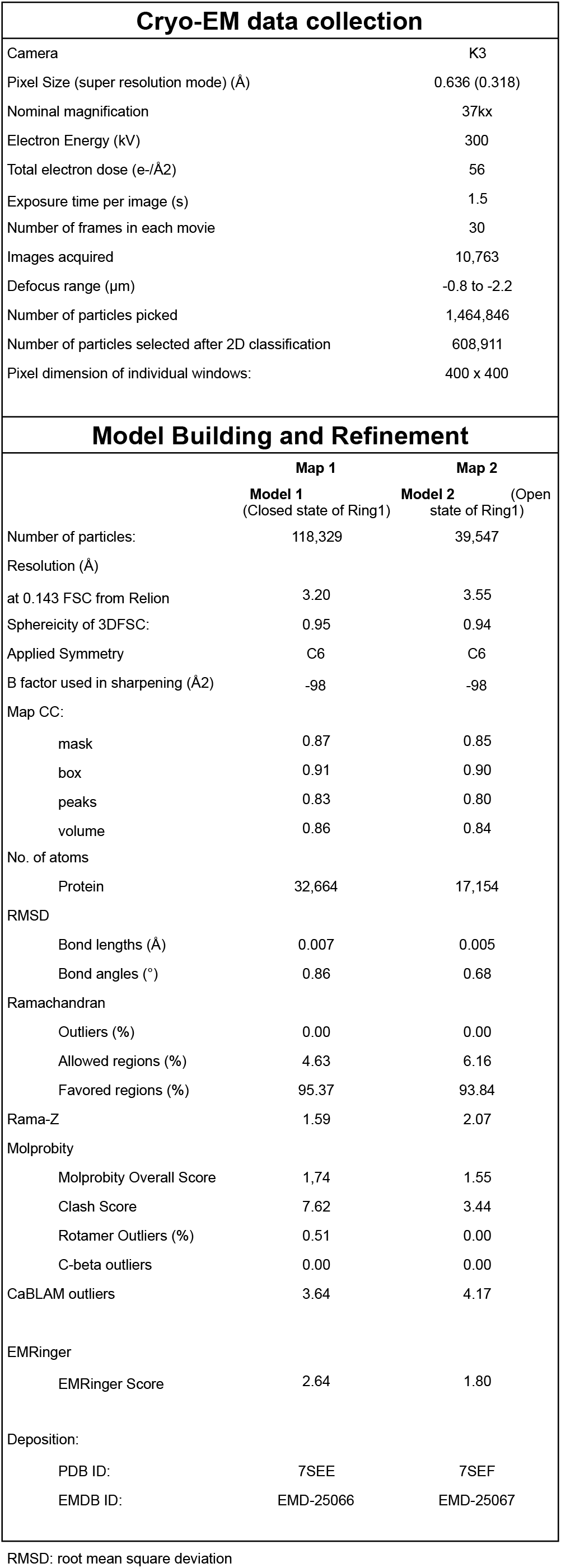
Data collection and refinement statistics for cryo EM structure of ΔRing 6, related to Figure 3

**Table S2:** Recombinant DNA used in this study

| Plasmid Name | Description | Reference | Addgene ID |
| --- | --- | --- | --- |
| pET17b-letAB / LetAB: WT | LetAB for complementation | Isom et al., 2017 | 175804 |
| pBEL1284 | His-LetA-LetB for expression | This study | 175960 |
| pBEL2004 | LetB(43-877; Δ634-746)-6xHis:Expression plasmid for 6 ring LetB deletion (without TM region) | Isom et al., 2020 | 139889 |
| pBEL2114 | His-LetA-LetB(Δ634-746) for expression | This study | 175961 |
| pBEL2116 | LetA-LetB(Δ48-159) for complementation | This study | 170393 |
| pBEL2117 | LetA-LetB(Δ160-279) for complementation | This study | 170394 |
| pBEL2118 | LetA-LetB(Δ280-391) for complementation | This study | 170395 |
| pBEL2119 | LetA-LetB(Δ392-515) for complementation | This study | 170397 |
| pBEL2120 | LetA-LetB(Δ516-633) for complementation | This study | 170398 |
| pBEL2121 | LetA-LetB(Δ634-746) for complementation | This study | 170399 |
| pBEL2122 | LetA-LetB(Δ747-877) for complementation | This study | 170399 |
| pBEL2196 | His-LetA-LetB(Δ702-717, replace with 114-130) for expression | This study | 170400 |
| pBEL2197 | His-LetA-LetB(Δ702-716, replace with 347-360) for expression | This study | 170401 |
| pBEL2198 | His-LetA-LetB(Δ702-716, replace with 580-600) for expression | This study | 170402 |
| pBEL2251 | His-LetA-LetB(Δ700-717, replace with 225-249) for expression | This study | 170403 |
| pBEL2252 | His-LetA-LetB(Δ700-716, replace with 445-479) for expression | This study | 170404 |
| pBEL2253 | His-LetA-LetB(Δ700-717, replace with 812-836) for expression | This study | 170405 |
| pBEL2265 | His-LetA-LetB(Δ347-360, replace with 702-716) for expression | This study | 175962 |
| pBEL2199 | LetA-LetB(Δ702-717, replace with 114-130) for complementation | This study | 170406 |
| pBEL2200 | LetA-LetB(Δ702-716, replace with 347-360) for complementation | This study | 170407 |
| pBEL2201 | LetA-LetB(Δ702-716, replace with 580-600) for complementation | This study | 170408 |
| pBEL2254 | LetA-LetB(Δ700-717, replace with 225-249) for complementation | This study | 170409 |
| pBEL2255 | LetA-LetB(Δ700-716, replace with 445-479) for complementation | This study | 170410 |
| pBEL2256 | LetA-LetB(Δ700-717, replace with 812-836) for complementation | This study | 170411 |
| pBEL2268 | LetA-LetB(Δ347-360, replace with 702-716) for complementation | This study | 175963 |
| pBEL2409 | LetA-[LetB(Δ711-715, replace with 355-359) for complementation | This study | 175964 |
| pBEL2412 | LetA-[LetB(Δ701-710, replace with 347-354) for complementation | This study | 175965 |
| pBEL2414 | LetA-LetB(P701N) for complementation | This study | 175966 |
| pBEL2415 | LetA-LetB(I703N) for complementation | This study | 175967 |
| pBEL2416 | LetA-LetB(A705N) for complementation | This study | 175968 |
| pBEL2417 | LetA-LetB(A706N) for complementation | This study | 175969 |
| pBEL2418 | LetA-LetB(G707N) for complementation | This study | 175970 |
| pBEL2419 | LetA-LetB(V708N) for complementation | This study | 175971 |
| pBEL2420 | LetA-LetB(P717N) for complementation | This study | 175972 |
| pBEL1863 | LetA-LetB(A473N) for complementation | This study | 170412 |
| pBEL1864 | LetA-LetB(W476N) for complementation | This study | 170413 |
| pBEL1865 | LetA-LetB(I477N) for complementation | This study | 170414 |
| pBEL1866 | LetA-LetB(L595N) for complementation | This study | 170415 |
| pBEL1867 | LetA-LetB(A598N) for complementation | This study | 170416 |
| pBEL1868 | LetA-LetB(L599N) for complementation | This study | 170417 |
| pBEL1869 | LetA-LetB(L711N) for complementation | This study | 170418 |
| pBEL1870 | LetA-LetB(I714N) for complementation | This study | 170419 |
| pBEL1871 | LetA-LetB(L715N) for complementation | This study | 170420 |
| pBEL1872 | LetA-LetB(F830N) for complementation | This study | 170421 |
| pBEL1873 | LetA-LetB(F833N) for complementation | This study | 170422 |
| pBEL1874 | LetA-LetB(I834N) for complementation | This study | 170423 |
| pBEL2343 | LetA-LetB(L123N) for complementation | This study | 170424 |
| pBEL2344 | LetA-LetB(L125N) for complementation | This study | 170425 |
| pBEL2345 | LetA-LetB(V126N) for complementation | This study | 170426 |
| pBEL1857 | LetA-LetB(L243N) for complementation | Isom et al., 2020 | 139904 |
| pBEL1858 | LetA-LetB(L246N) for complementation | Isom et al., 2020 | 139905 |
| pBEL1859 | LetA-LetB(V247N) for complementation | Isom et al., 2020 | 139906 |
| pBEL1860 | LetA-LetB(L355N) for complementation | Isom et al., 2020 | 139907 |
| pBEL1861 | LetA-LetB(L358N) for complementation | Isom et al., 2020 | 139908 |
| pBEL1862 | LetA-LetB(L359N) for complementation | Isom et al., 2020 | 139909 |

